# Genetic adaptation to pathogens and increased risk of inflammatory disorders in post-Neolithic Europe

**DOI:** 10.1101/2022.07.02.498543

**Authors:** Gaspard Kerner, Anna-Lena Neehus, Laurent Abel, Jean-Laurent Casanova, Etienne Patin, Guillaume Laval, Lluis Quintana-Murci

**Affiliations:** Institut Pasteur, Université Paris Cité, CNRS UMR2000, Human Evolutionary Genetics Unit, F-75015 Paris, France; Laboratory of Human Genetics of Infectious Diseases, INSERM UMR 1163, Necker Hospital for Sick Children, Paris, France; University Paris Cité, Imagine Institute, Paris, France; St. Giles Laboratory of Human Genetics of Infectious Diseases, The Rockefeller University, New York, NY, United States; Howard Hughes Medical Institute, New York, NY, United States; Department of Pediatrics, Necker Hospital for Sick Children, Paris, France; Collège de France, Chair of Human Genomics and Evolution, F-75005 Paris, France

**Keywords:** ancient DNA, immunity, host defense, natural selection, local adaptation, inflammatory disorders, approximate Bayesian computation, antagonistic pleiotropy, *LBP*

## Abstract

Ancient genomics can directly detect human genetic adaptation to environmental cues. However, it remains unclear how pathogens have exerted selective pressures on human genome diversity across different epochs and affected present-day inflammatory disease risk. Here, we use an ancestry-aware approximate Bayesian computation framework to estimate the nature, strength, and time of onset of selection acting on 2,879 ancient and modern European genomes from the last 10,000 years. We found that the bulk of genetic adaptation occurred after the start of the Bronze Age, <4,500 years ago, and was enriched in genes relating to host-pathogen interactions. Furthermore, we detected directional selection acting on specific leukocytic lineages and experimentally demonstrated that the strongest negatively selected immunity gene variant — the lipopolysaccharide-binding protein gene (*LBP*) D283G — is hypomorphic. Finally, our analyses suggest that the risk of inflammatory disorders has progressively increased in post-Neolithic Europeans, partly due to antagonistic pleiotropy following genetic adaptation to pathogens.

## INTRODUCTION

Infectious diseases have been the leading cause of human mortality throughout human history.^1,2^ Population genetics studies have provided support for the notion that pathogens are among the strongest selective forces faced by humans since the discovery in the 1950s that heterozygosity for red blood-cell disorders provides some protection against malaria.^3^ An increasing number of genes involved in host-pathogen interactions have since been identified as targets of natural selection.^4–8^ However, major questions remain regarding the evolutionary impact of infectious diseases on human genome diversity. First, little is known about the specific epochs during which humans were most exposed to pathogens and pathogen-mediated selection. It has been suggested that the transition to an agriculture-based lifestyle, which began ~10,000 years ago, increased exposure to deadly microbes, including density-dependent viruses and zoonoses,^9–11^ but the archaeological and genetic evidence is scarce, and even challenges this view.^12,13^ Second, the extent to which host defenses, including the engagement of leukocytic lineages, the qualitative and quantitative composition of which is associated with common and rare disorders,^14–17^ have been affected by such exposure has not been explored. Third, the rising life expectancy in recent centuries^18^ has contributed to an increase in the prevalence of inflammatory and autoimmune disorders, but it has been hypothesized that this increase is also the result of long-term pathogen pressures and antagonistic pleiotropy of the selected gene products.^19–21^

The antagonistic pleiotropy hypothesis is supported by the overlap between loci underlying infectious and inflammatory traits^22,23^ and the discovery of several pleiotropic variants conferring protection against infectious diseases and susceptibility to some chronic inflammatory or auto-immune conditions.^22–27^ Classic examples are the human leukocyte antigen (*HLA*) loci, variants of which are thought to be under pathogen-driven positive selection,^28^ and to increase considerably the risk of autoimmune disease (e.g. *HLA-B27* and spondyloarthritis and *HLA-DQ8* and diabetes).^29–33^ Another example is the common *TYK2* P1104A variant, which protects against auto-immune phenotypes (OR: 0.1-0.3) but, in the homozygous state, confers a predisposition to mycobacterium-related infectious diseases (OR >10).^25,26,34–39^ Nevertheless, the evidence that antagonistically pleiotropic variants have been selected in humans remains circumstantial.

Detection of a legacy of positive selection (rapid increase in the frequency of a beneficial allele) or negative selection (the removal of a deleterious variant) in human populations has long been limited to statistical inference from patterns of genetic variation in contemporary individuals. However, in recent years, the increasing availability of ancient DNA (aDNA) data has greatly facilitated the study of human genetic adaptation over time. Direct comparisons of past and current allele frequencies have provided new insight into the genes and functions involved in human adaptation to environmental changes following cultural transitions.^40–42^ A pioneering study based on 230 Eurasian samples across the Holocene period detected 12 positively selected loci associated with diet and skin pigmentation, but also host defense against pathogens.^43^ Another study explored how the arrival of Europeans in the Americas altered the exposure of Native Americans to new pathogens. Comparisons of genomic data from a Canadian First Nation population dating from before and after the first contact with Europeans showed a recent decrease in the frequency of formerly beneficial *HLA-DQA1* alleles in the native population.^44^ These studies highlight how the analysis of ancient genomes can identify specific genetic variants under selection, but we still lack a comprehensive picture of the selective forces affecting host defense genes during human history and of the times at which these forces operated.

Ancient genomics provides us with a unique opportunity to determine whether resistance to infectious diseases and susceptibility to inflammatory diseases have changed in the recent past. Furthermore, aDNA data can be used to detect variants subject to complex models of selection, such as time-dependent selection, including selection on *de novo* or standing variation due to sudden environmental changes (e.g., epidemics). For example, a recent study revealed that negative selection has led to a rapid decrease in the frequency of a 30,000-year-old tuberculosis (TB) risk variant over the last 2,000 years, suggesting that TB has recently imposed a heavy burden in Europeans.^35^ In this study, we sought to reconstruct the history of host-pathogen interactions by detecting immunity variants affected by natural selection, thereby modulating infectious and/or inflammatory disease risk, over the last 10,000 years. To this end, we explored the strength and timing of both positive and negative selection at the genome-wide scale, by analyzing 2,879 ancient and modern European genomes in an ancestry-aware approximate Bayesian computation (ABC) framework.

## RESULTS

### Inferring the intensity and onset of positive selection from ancient DNA data

We assembled a genome-wide dataset corresponding to 2,376 ancient and 503 modern individuals of western Eurasian ancestry, and computed allele frequency trajectories for 1,233,013 polymorphic sites, over a time transect covering the Neolithic, the Bronze Age, the Iron Age, the Middle Ages and the present (**Methods**). Using simulation-based ABC^45^ and the calculated trajectories, we estimated the selection coefficient (*s*) and the time of selection onset (*T*) for each derived allele. We considered a variant to be a candidate for positive selection if its *s* value was higher than that of 99% of *s* estimates for simulated neutral variants (*p*_sel_ <0.01). We used a demographic model^35^ accounting for (i) the major migratory movements contributing to the genetic diversity of contemporary Europeans, i.e., the arrival of Anatolian farmers ~8,500 years ago (ya)^46,47^ and that of populations associated with the Yamnaya culture around ~4,500 ya,^48,49^ (ii) the uneven sampling across time, with the matching of simulated data on the observed numbers of ancient and modern DNA data, and (iii) ancestry variation across epochs, by matching on observed Anatolian and Pontic Steppe ancestries (**Methods**).

Our simulations showed that the estimation of *s* was highly accurate (*r^2^* = 0.93 between simulated and estimated values; **Figure S1A**), and the power to detect selection from *s* was >80% for variants with a present-day frequency >20%, regardless of the timing of selection onset (**Figure S2**). The accuracy of estimation was lower for *T*(*r*^2^ = 0.76; **Figure S1B**), but our approach was nevertheless able to distinguish between variants selected before and after the beginning of the Bronze Age, ~4,500 ya (F1 score = 0.85; **Figure S1C**). The application of ABC to the observed allele frequency trajectories replicated the 12 loci previously shown to be subject to positive selection in Europe^43^ (based on the same criterion of ≥3 candidate variants for positive selection per locus). These loci included genes associated with host defense (*HLA* and *TLR10*-*TLR1*-*TLR6*), skin pigmentation (*SLC45A2* and *GRM5*) and metabolism (*MCM6/LCT, SLC22A4* and *FADS1*) (**Figure 1; Table S1**). As further evidence of the accuracy of our approach, the *s* estimate for the lactase persistence allele (rs4988235) was 8.1% (95% CI: [0.06-0.09]) with selection beginning 6,102 ya (3,150-8,683), as previously reported.^50,51^ Our approach expands previous aDNA studies by not only identifying known and novel candidates for selection, but also providing appropriate estimates of the selection parameters, *s* and *T*, over the entire genome.

**Figure 1.**
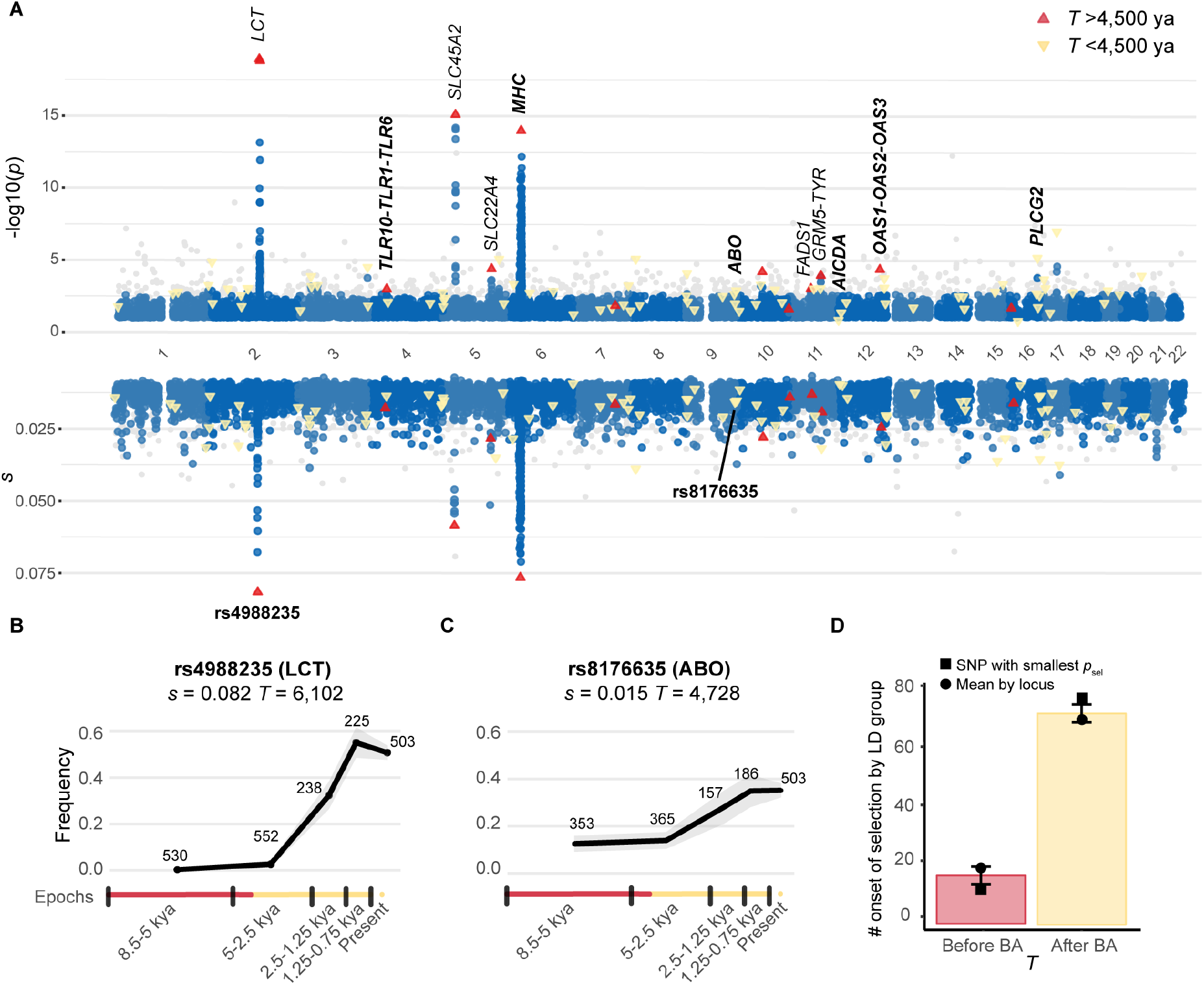
Genome-wide detection of positive selection in Europe since the Neolithic. (A) Test for positive selection (−log10(*p*_sel_)) and selection coefficient estimates (*s*) for each genomic marker in the capture dataset with *p*_sel_ < 0.1. The empirical null distribution was approximated by a beta distribution (**Methods**) Variants with *p*_sel_ < 10^−4^ outside the 89 enriched loci are colored in gray. Several candidate genes for positive selection are indicated, and host defense genes are highlighted in bold. Upward-pointing pink triangles and downward-pointing yellow triangles correspond to the 89 SNPs with the smallest *p*_sel_ at each of the 89 candidate loci selected before and after the beginning of the Bronze Age, respectively. The time of selection onset was estimated as the mean *T* across all SNPs with *p*_sel_ <0.01 at each locus. (B-C) Frequency trajectory of the most significant variant at (B) the *LCT* locus (rs4988235) and (C) the *ABO* locus (rs8176635). Gray shading around the frequency trajectory indicates the lower and upper bounds of the 95% CI for variant frequency estimation in each epoch. (D) Mean number of times a single randomly selected candidate SNP per LD group had an estimated *T* before or after the beginning of the Bronze Age, across 1,000 replicates. The error bar for each bar plot indicates the standard deviation of the distribution obtained. We also included the estimate based on the mean *T* of all SNPs with *p*_sel_ <0.01 (as for the Manhattan plot) for each of the 89 LD groups (rectangles), or based on the SNP with the smallest *p*_sel_ for each LD group (circles).

### Positive selection has pervasively affected host defense genes in Europeans

We identified the strongest signals of time-dependent positive selection, by focusing on the top 2.5% of loci with the highest proportions of candidate variants, i.e., regions defined on the basis of linkage disequilibrium (LD) groups containing >7.5% of the candidate positively selected, derived variants (*p*_sel_ <0.01, **Methods**). We identified 102 candidate loci for positive selection, 89 of which were non-consecutive. These loci were enriched in gene ontology (GO) categories such as *antigen processing and presentation of endogenous antigen* (Bonferroni corrected *p*_adj_ = 0.01), *viral life cycle* (*p*_adj_ = 0.03), and *positive regulation of leukocyte activation* (*p*_adj_ = 0.05), as well as *transport vesicle, vesicle membrane, luminal side of endoplasmic reticulum membrane, cell surface* and *vesicle-mediated transport*. These 89 loci were also enriched in a curated list of immunity genes (IGs; **Methods**), whether we considered all candidate genes (OR = 1.6, *p* = 8.0 × 10^−3^) or a single gene per locus (28/89 loci; OR = 1.6, *p* = 0.049). These genes included the *OAS* cluster (*OAS1-OAS2-OAS3-RNASEL*), which modulates the response to infection with RNA viruses,^52^ and genes underlying inborn errors of immunity (IEI), such as *AICDA*, for which biallelic loss-of-function mutations cause antibody deficiencies (OMIM:6055258 and OMIM:605257), and *PLCG2*, for which monoallelic gain-of-function mutations underlie autoinflammatory disorders (OMIM:614878 and OMIM:614468).

One of the strongest signals identified was close the *ABO* gene (smallest *p*_sel_, **Table S2**), which encodes the ABO blood group system. We found that alleles tagging the A and B blood groups^53^ were candidates for positive selection (*p*_sel_ <0.01; **Table S3**), suggesting that the frequency of individuals with the A, AB and, particularly, B blood groups has increased through positive selection over the last few millennia. The A and B blood groups have been shown to confer limited protection (OR >0.94) against childhood ear infections,^54,55^ and mild susceptibility (OR <1.13) to malaria and COVID-19,^53,56^ consistent with the action of balancing selection.^57^ Our results show that the bulk of recent genetic adaptation in Europe has primarily concerned genes involved in the host response to infection, suggesting that pathogens have imposed strong selective pressures over the last few millennia.

### Genetic adaptation has occurred principally since the Neolithic period

We then explored the time at which positive selection began at the 89 loci, by determining whether selection occurred before or after the start of the Bronze Age (**Figure S1C**). For more than 80% of loci, selection was estimated to have begun after the beginning of the Bronze Age (<4,500 ya) (**Figure 1**). Indeed, the distribution of *T* estimates for the top 89 variants was not consistent with a uniform distribution across time (*p* = 1.3 × 10^−4^), with a much more recent time of selection onset than expected (*T*_mean_ = 3,327 ya; **Figure S3A; Table S2**). This result cannot be explained by differences in detection power, as our approach has a higher power for variants with selection beginning at earlier timepoints (**Figure S2**). Furthermore, using 100 distributions of *T* estimates obtained from 89 simulated variants drawn with uniform durations of selection and matched for allele frequency and selection intensity, we checked that the distributions were not skewed towards *T* estimates postdating the beginning of the Bronze Age (*p*_adj_ > 0.05 for all 100 distributions; **Methods**). Finally, the ancient and modern DNA datasets were processed differently, so batch effects could have resulted in a sudden change in allele frequency in the last epoch and, therefore, a bias of *T* towards more recent times. However, when estimating *T* for aDNA only, we again obtained a significantly larger fraction of estimates < 4,500 ya (*n* = 67/89;*p* < 10^−3^; **Figure S3B**). These analyses support a scenario in which most of the positive selection events have occurred since the Neolithic Period.

In light of these results, we reasoned that the arrival of populations from the Pontic-Caspian region ~4,500 ya may have brought beneficial variants that continued to be subject to positive selection in the resulting admixed population, a process known as adaptive admixture.^58^ In this case, we would expect to see a positive correlation between Pontic Steppe ancestry and the probability of carrying positively selected alleles. However, no significant differences were observed between positively selected alleles and matched controls (*p*_adj_ >0.05 for all matched samples; **Methods**), and there was no systematic bias in the ancestry of simulated and observed aDNA samples (**Figure S4A**). Our results therefore rule out adaptive admixture as the main driver of positive selection in post-Neolithic Europe, instead suggesting that there has been selection on standing variation; in other words, most of the alleles that became beneficial after the Bronze Age were already present in Neolithic Europe (**Figure S4B**).

### Regulatory elements of immunity genes have been a prime substrate of positive selection

We then investigated *in-silico* the functional effects of candidate positively selected variants (*n* = 1,846). We found that these variants were strongly enriched in missense variants (OR = 3.8, *p* = 8.8 × 10^−21^) (**Figure 2A**). Furthermore, IGs harboring missense variants were found to be significantly enriched in positively selected variants relative to the rest of the candidate loci (*p* = 6.0 × 10^−3^). At the genome-wide scale, we detected 11 missense variants of IGs with signals of positive selection (*MYH9, CHAT, NLRC5, INPP5D, SELE, C3, PSMB8, CFB, PIK3R1, RIPK2, CD300E*). Eight of these variants were associated with immune cell traits and infectious or autoimmune disorders, such as blood cell counts, viral warts and type I diabetes, in phenome-wide association studies (*p* < 10^−5^; **Figure 2B**). However, given that most positively selected variants (>96%) are non-coding, we investigated whether variations of gene expression could account for the observed selection signals. Taking LD and derived allele frequency (DAF) (*r*^2^ < 0.6) into account, we found that candidate variants were enriched in *cis*-eQTLs in whole blood, particularly for strong eQTL associations (e.g., OR = 2.9;*p* = 3.3 × 10^−21^ for eQTLs with *p* <10^−50^; **Figure 2A**; **Methods**). For example, the top candidate variant for the *OAS* cluster, rs1859332>T, was found to be linked (*r^2^* = 0.97) to the *OAS1* splice QTL rs10774679>C (isoform p46), which is of Neanderthal origin.^59^ Interestingly, this isoform has been associated with protection against severe COVID-19 (OR = 0.05),^60,61^ suggesting that past coronavirus-like viruses may have driven selection at this locus.

**Figure 2.**
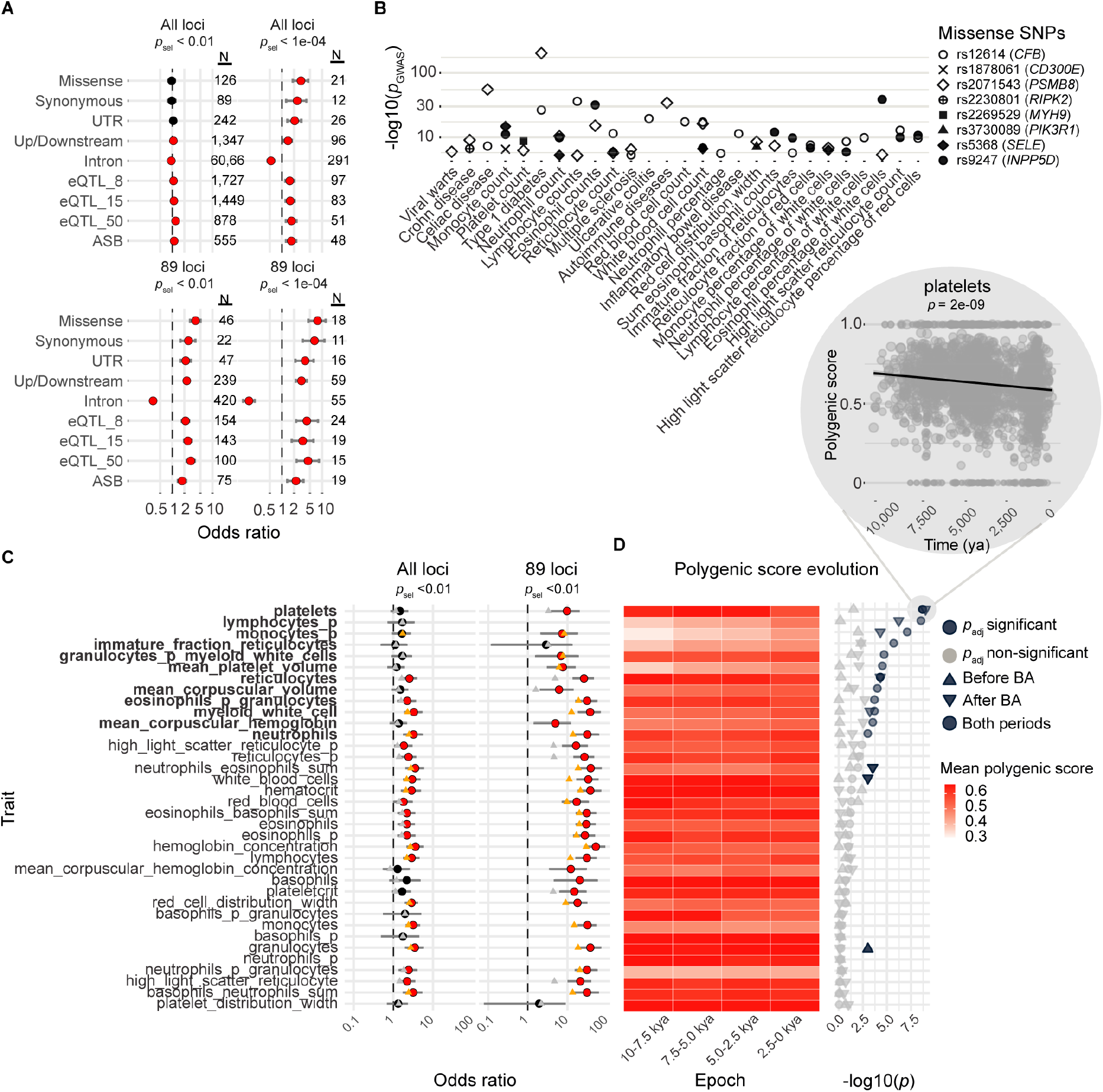
Genetic variants and host defense traits targeted by positive selection. (A) Enrichments in missense, whole blood *cis*-eQTL or ASB variants for variants with *p*_sel_ <10^−2^ and *p*_sel_ <10^−4^, across the genome (‘All loci’) or within the 89 candidate positively selected loci (‘89 loci’). Only variants of transcripts annotated as ‘protein-coding’ were considered for the analysis of variant annotations. The three eQTL groups were defined on the basis of the statistical significance of the eQTL association, *p*_eQTL_ < 5 × 10^−8^, <10^−15^ or <10^−50^. (B) Significant associations (*p* <10^−5^; https://genetics.opentargets.org) in PheWAS between positively selected missense variants overlapping host defense genes and hematopoietic traits, infections or inflammatory disorders. (C) Enrichments in variants associated with the 36 hematopoietic traits studied for variants with *p*_sel_ <10^−2^, across the genome (‘All loci’) or within the 89 candidate positively selected loci (‘89 loci’). Red and black circles indicate significant and non-significant enrichments, respectively. Orange and gray triangles indicate significant and non-significant enrichments, respectively, after exclusion of the *HLA* locus. (D) Left: mean polygenic score (**Methods**) across four equally spaced epochs for each of the hematopoietic traits. Right: −log10 *p* values for the significant increases or decreases in each hematopoietic trait over the last 10,000 years (dot), before (upward-pointing triangle) or after (downward-pointing triangle) the beginning of the Bronze Age. Light gray symbols represent non-significant trajectories. The inset shows the decrease in polygenic score over time for platelet counts, which has the highest −log10 *p* value. The black line is the regression line. (CD) Traits followed by ‘_p’ indicate the percentage of the cell type considered, whereas other traits are absolute counts.

Positively selected variants are also enriched in variants affecting transcription factor (TF) binding (OR = 1.75; *p* = 1.6 × 10^−6^; **Figure 2A**), based on allele-specific binding (ASB) events detected in ChIP-Seq data covering 1,025 human TFs and 566 cell types (FDR < 0.01)^62^. For example, the positively selected rs3771180>T variant (*s* = 0.015, *T* = 2,250, *p*_sel_ = 0.005) at the *IL1RL1* locus disturbs the 7 bp binding motif of *JUND* (OR = 3.07;*p*_adj_ = 5.7 × 10^−12^) and is associated with a lower risk of asthma^63^ and higher neutrophil levels.^14^ Moreover, 17 of the 265 TFs with at least five ASBs have ASB-associated variants enriched in positive selection signals, and the genes closest to these ASBs are enriched in IGs (OR = 1.24; *p* = 0.04). Collectively, these results provide insight into the regulatory mechanisms underlying recent human adaptation in the context of immunity to infection.

### Directional selection has affected leukocytic lineages since the Bronze Age

We analyzed epigenomic data from ENCODE,^64^ to identify the tissues for which the regulatory elements were the most enriched in positively selected variants. In tests with matched controls and correction for multiple testing, we found an enrichment in positively selected variants at DNase-hypersensitive sites in 24 of the 41 tissues and cell types tested, including monocytes (*p*_adj_ = 6.1 × 10^−7^), tonsils (*p*_adj_ = 2.1 × 10^−6^) and blood (*p*_adj_ = 1.0 × 10^−5^) (**Methods**). These results, together with the enrichment in ASB in candidate variants close to IGs, led us to investigate whether positively selected variants affect hematopoiesis by altering blood cell fractions.

Taking LD and DAF into account, we found that variants associated with blood cell composition in GWAS were strongly enriched in positively selected variants (OR = 10, *p* = 7.2 × 10^−65^) (**Figure 2C**). For each of the 36 hematopoietic traits, we analyzed the frequency trajectories over time of all the trait-increasing alleles (*p*_GWAS_ < 5.0 × 10^−8^) with a polygenic score derived from GWAS data,^40^ within the studied candidate positively selected LD groups (**Figure S5; Methods**). This score integrates tens or hundreds of generally small effect sizes, which we assessed over hundreds of generations. We found that polygenic scores for platelet and reticulocyte counts had decreased significantly over the last 10,000 years, whereas scores for mean platelet volume and mean corpuscular (red blood cell) volume, and the mean mass of hemoglobin per red blood cell, had significantly increased (**Figure 2D**). Likewise, polygenic scores for the proportion of eosinophils among granulocytes were found to have significantly decreased over the study period, whereas the proportions of lymphocytes and monocytes among white blood cells and neutrophil counts had significantly increased (**Figure 2D**). Importantly, the polygenic scores of seven traits, including traits relating to platelets, reticulocytes, lymphocytes and monocytes, were found to have significantly increased or decreased in post-Neolithic Europe, whereas no trait other than the number of granulocytes changed significantly during the Neolithic (**Figure 2D**), consistent with directional selection acting principally on hematopoietic lineages after the Bronze Age.

### Joint temporal increase in resistance to infection and the risk of inflammation

It has been suggested that past selection relating to host-pathogen interactions has favored host resistance alleles that, today, increase the prevalence of chronic inflammatory disorders due to antagonistic pleiotropy.^4,19–21,23,24^ We investigated the correlation of genetic susceptibility to inflammatory disorders with resistance to infectious disease, by assembling the results for (i) 40 GWAS of infectious diseases, including severe infections, such as TB, hepatitis B and C, AIDS and COVID-19 (COVID-19 HGI, release 7), and surrogate phenotypes, such as *positive TB test* (based on tuberculin skin test) or *tonsillectomy*, and (ii) 30 GWAS of inflammatory/auto-immune disorders, including rheumatoid arthritis (RA), systemic lupus erythematosus (SLE) and inflammatory bowel disease (IBD) (**Table S4**). For both independent and randomly matched variants, we found that variants significantly associated with both infectious and inflammatory traits (i.e., pleiotropic variants; *p*_GWAS_ < 5.0 × 10^−8^) were more prevalent than expected by chance (OR = 132, *p* = 7.4 × 10^−28^ and *p* <10^−3^, respectively; **Methods**), suggesting a shared genetic architecture. Furthermore, that the estimated *s* values for these pleiotropic variants are stronger than for randomly matched variants (Wilcoxon *p* = 0.02) supports their adaptive nature.

We used polygenic risk scores (PRS) to explore changes in the frequency of risk alleles for infectious or inflammatory disorders (*p*_GWAS_ <5 × 10^−8^) over time (**Methods**). We found that the PRS for the merged set of all inflammatory/autoimmune disorders had significantly increased over the last 10,000 years, whereas that for infectious diseases had significantly decreased (**Figure 3A**). Taking into account the coverage, location and ancestry of the samples, the risk of Crohn’s disease (CD) or of IBD generally has increased significantly, whereas that of severe COVID-19 has decreased significantly, essentially since the Neolithic period (**Figure 3A-B**). Given the recent nature of the emergence of SARS-CoV-2, the infectious agent of COVID-19, this finding suggests that other related pathogens have exerted selection pressure in recent European history.

**Figure 3.**
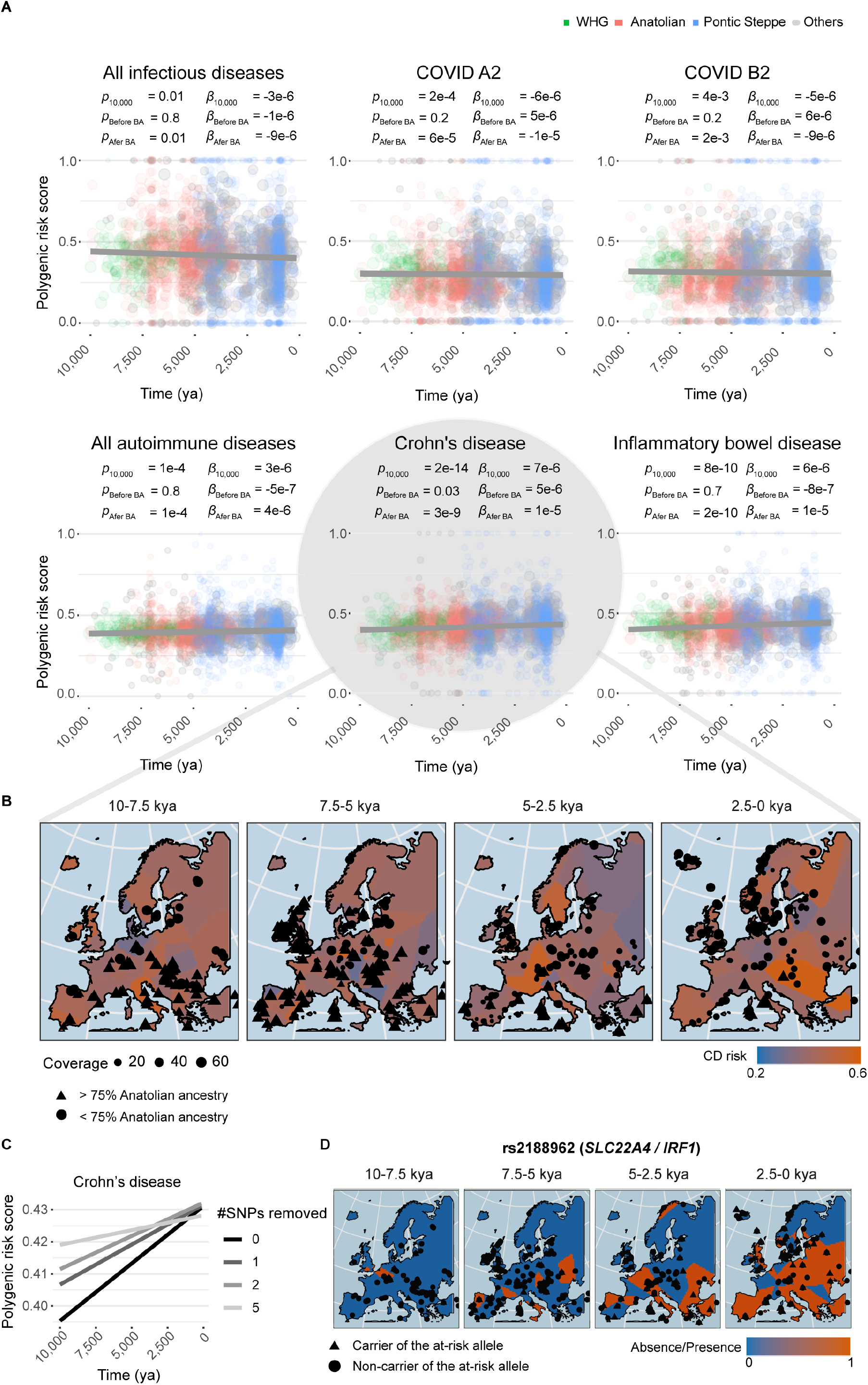
Resistance to infection and risk of inflammation have increased since the Neolithic. (A) Polygenic scores for infectious and autoimmune traits as a function of time, over the last 10,000 years. COVID A2 and B2 indicate critical COVID-19 cases vs. the general population and hospitalized COVID-19 cases vs. the general population, respectively. Dot size scale with the number of SNPs genotyped in the individual. Dark gray lines are the regression lines for a model adjusted for ancestry and geographic location. *P*-values indicate the significance of the regression model, over the last 10 millennia (*p*), before the Bronze Age (*p*_Before BA_) or since the beginning of the Bronze Age (*p*_After BA_) (**Methods**). Beta values for the regression model considering only samples dating from before (β_Before BA_) or after (β_After BA_) the beginning of the Bronze Age are shown. Green, pink and blue dots indicate individuals with >75% Western hunter-gatherer, Anatolian or Pontic Steppe ancestry, respectively. Individuals with mixed ancestries are shown in gray. (B) Polygenic score for Crohn’s disease as a function of the geographic location and age of the samples, obtained with the “bleiglas” package of R version 3.6.2. Individuals with >75% Anatolian ancestry are represented by triangles, and the others are represented by circles. (C) Polygenic risk score for Crohn’s disease as a function of time, after removal of the SNPs most significantly associated with sample age. (D) Presence or absence of the CD-risk allele rs2188962>T as a function of the geographic location and the age of ancient samples.

We then searched for the specific genetic variants making the largest contribution to the changes over time in the risks of IBD and severe COVID-19. Taking multiple testing into account, we found that four risk alleles for IBD and one protective allele for COVID-19 had significantly increased in frequency over time (**Figure 3C** and **Table S5**). The risk alleles concerned were the IBD- and/or CD-associated rs1456896>T, rs2188962>T, rs11066188>A and rs492602>G alleles, and the COVID-19-associated rs10774679>C allele. The first three of these variants increase the expression of *IRF1, SH2B3*, and *IKZF1*, respectively, in blood (rs2188962, *s* = 0.022, *T* = 8,075; rs11066188, *s* = 0.016, *T* = 4,045; rs1456896>C, *s* = 0.007, *T* = 3,264; **Figure 3D**). These genes have been shown to protect against several infectious agents,^27,65,66^ suggesting that increases in their expression may be beneficial, even if they slightly increase autoimmunity risk. The fourth IBD-associated risk allele, rs492602>G (*s* = 0.0015, *T* = 6,823), is a *FUT2* variant in complete LD (*r*^2^ = 0.99) with the null allele rs601338>A, which confers monogenic resistance to infections with intestinal norovirus^67^ and respiratory viruses.^68^ Finally, the COVID-19-associated variant rs10774679>C (*s* = 0.013, *T* = 3,617) is linked (*r*^2^ = 0.9) to the *OAS1* splice variant rs10774671>G, which is also associated with slightly higher susceptibility to IBD (OR = 1.08; *p* = 1.4 × 10^−3^; FINNGEN_R5_K11_IBD). Overall, these analyses provide support for a role of selection in increasing autoimmune disease risk over recent millennia, particularly for gastrointestinal inflammatory traits, probably due to antagonistic pleiotropy.

### Searching for the footprints of time-dependent negative selection

Given the observed links between candidate selected variants and host defense, we then used our approach to identify candidate variants increasing infectious disease risk. Specifically, we searched for variants of genes involved in host-pathogen interactions that have been under time-dependent negative selection, that is, variants that have become deleterious since the Neolithic. A typical example is *TYK2* P1104A, which we have previously shown to underlie clinical TB^34,39^ and to have evolved under negative selection over the last 2,000 years, probably due to an increase in the pressure imposed by *Mycobacterium tuberculosis*.^35^ We removed the 89 positively selected candidate regions, and loci presenting positive selection signals in contemporary Europeans (**Methods**), and then searched for candidate deleterious variants displaying significant decreases in DAF in the rest of the genome (93% of all LD groups).

We first checked that the *s* and *T* estimates for alleles under negative selection were as accurate as those for positively selected alleles (**Figure S1D-E**). We confirmed that variants with a low DAF (<5% across epochs) were strongly enriched (OR = 4.8, *p* = 2.7 × 10^−210^) in candidate negatively selected variants, as opposed to positively selected alleles (OR = 0.64, *p* = 4.2 × 10^−7^). We found that negative selection had a broader impact across the genome; 25% of LD groups harbor candidate negatively selected variants (*p*_sel_ < 10^−4^), whereas only 3% harbor positively selected alleles (**Figures 1A** and **4A**). By focusing on negatively selected missense variants at conserved positions (GERP score >4; n = 50; **Table S6**), we found that the vast majority of these variants (41 out of the 50; 82%), like the positively selected variants, had an estimated onset of selection <4,500 ya (**Figure 4**). Focusing on immunity genes, we identified six negatively selected missense variants: *LBP* D283G (rs2232607), *TNRC6A* P788S (rs3803716), *C1S* R119H (rs12146727), *IL23R* R381Q (rs11209026), *TLR3* L412F (rs3775291) and *TYK2* P1104A (rs34536443). These time-series analyses provide compelling evidence for negative selection against variants of host defense genes in recent millennia.

**Figure 4.**
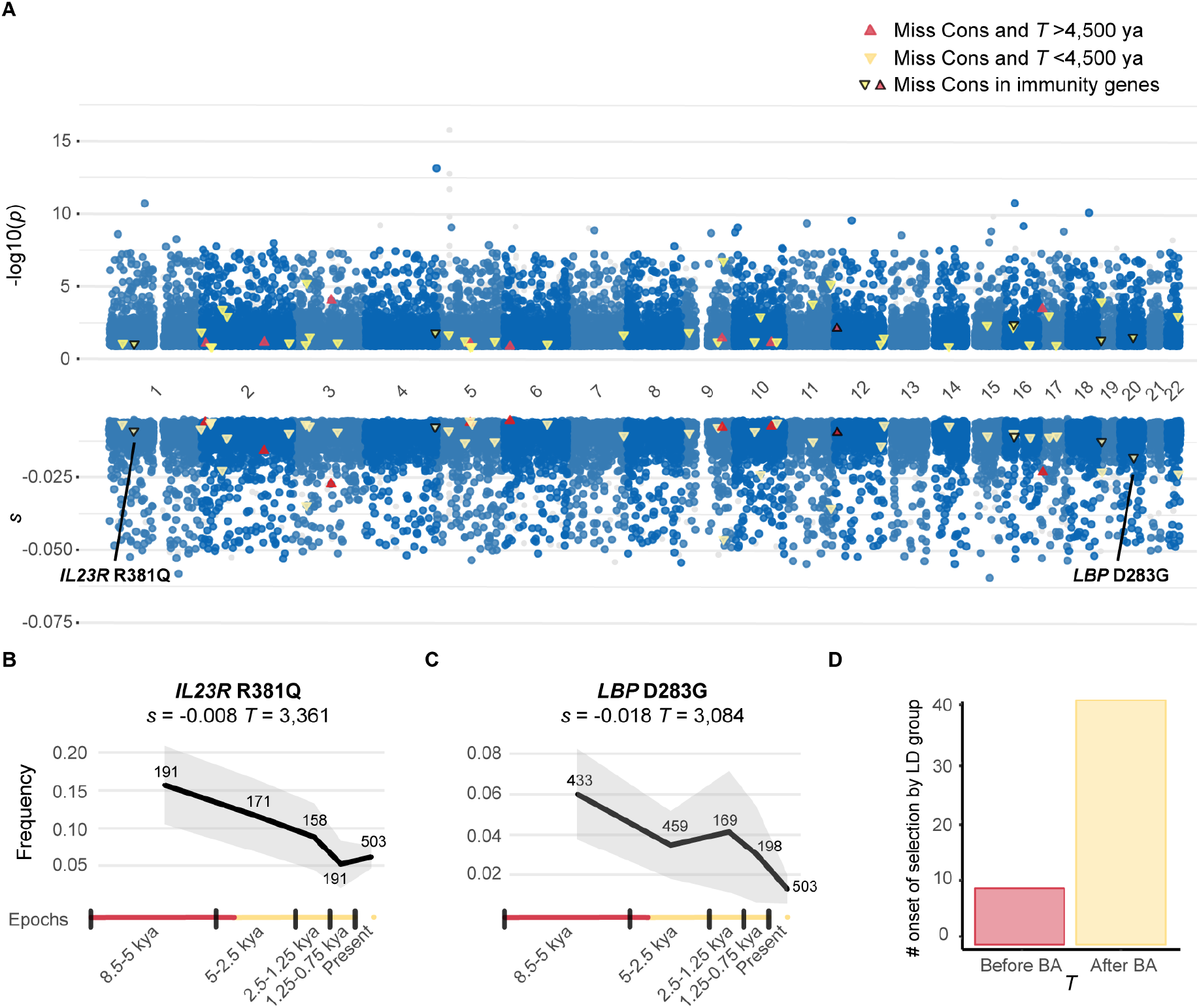
Genome-wide detection of negative selection since the Neolithic. (A) Test for negative selection (−log10(*p*)) and selection coefficient estimates (*s*) for each genomic marker in the capture dataset. Red and yellow triangles indicate missense variants at conserved positions (GERP score >4) under negative selection, starting before or after the beginning of the Bronze Age, respectively. The triangles for the six candidate negatively selected variants of immunity genes (IGs) are outlined in black. (B-C) Frequency trajectory for the strongest candidate negatively selected variant, *LBP* D283G, and the missense variant *IL23R* R381Q. (D) Number of candidate variants under negative selection beginning before or after the beginning of the Bronze Age.

### The D283G variant is hypomorphic and impairs LBP expression

We investigated the functional impact of negatively selected variants, by focusing on the *LBP* D283G variant, which had the strongest negative selection signal (*s* = −0.018, *T* = 3,084). *LBP* encodes the lipopolysaccharide (LPS)-binding protein, which senses LPS — a major component of the outer membrane of Gram-negative bacteria — and initiates immune responses that prime host defense mechanisms against further infection.^69^ Previous studies have shown that a common *LBP* variant (P333L; 8% in Europe) reducing both LPS binding capacity and cytokine response is associated with greater mortality due to sepsis and pneumonia.^70^ We investigated whether *LBP* D283G, which has decreased in frequency from 6% to 1.2% over the last 5,000 years, had a similar biochemical impact on LBP function, potentially decreasing host fitness for fighting bacterial infections.

We transiently transfected HEK 293T cells with plasmids encoding wild-type (WT) or mutant LBP cDNAs, including the previously reported P333L variant. All constructs were C-terminally tagged with DDK and equal transfection efficiencies were confirmed by RT-qPCR (**Figure 5A**). Protein levels were analyzed by western blot on whole-cell lysates and cell culture supernatants. The WT and mutant proteins were detected at the expected molecular weight (~60 kDa) with an LBP-specific antibody and an antibody against the C-terminal tag (**Figure 5B**). However, D283G levels were lower than WT protein levels, and P333L was not detectable. The low levels of D283G protein were confirmed by ELISA (**Figure 5C**), suggesting that this variant may be less stable when secreted. Finally, we showed that the D283G and WT proteins had similar LPS binding capacities (**Figure 5D**), demonstrating that this variant does not affect this function. These findings suggest that D283G alters LBP stability, resulting in lower LBP protein levels, potentially impairing the host response to bacterial exposure.

**Figure 5.**
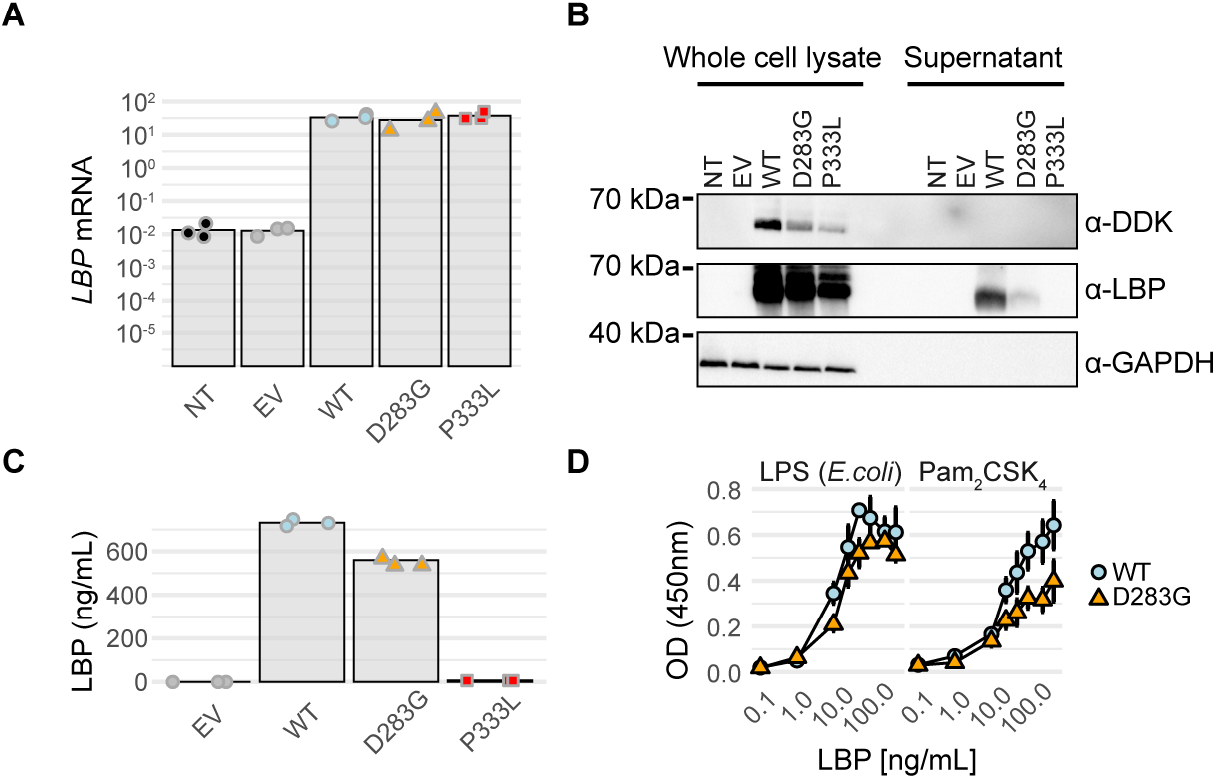
*LBP* D283G is hypomorphic in an overexpression system. (A) RT-qPCR for *LBP* on cDNA from HEK293T cells non-transfected (NT) or transfected with an empty plasmid (EV), or plasmids encoding wild-type (WT), or mutated *LBP*. Dots indicate three independent experiments and the height of each bar their mean values. (B) Western Blot of whole cell lysates or cell culture supernatants from HEK293T cells either left NT, transfected with an EV, or C-terminally tagged plasmids expressing WT or mutated *LBP*. LBP was detected with a polyclonal anti-LBP antibody and an antibody against the C-terminal DDK tag. An antibody against GADPH was used as loading control. The results shown are representative of three independent experiments. (C) LBP concentration in cell culture supernatants from transfected HEK293T cells as measured by ELISA. Dots indicate three independent experiments and the height of each bar their mean values. (D) Binding of WT or mutant LBP collected from cell culture supernatant at increasing concentrations to LPS and Pam2CSK4 assessed by a binding assay. Each point represents the mean of two biological replicates ±SD.

## DISCUSSION

Based on a time-series of human aDNA data and extensive computer simulations, we have delineated the genomic regions under the strongest selective pressure over the last 10,000 years of European history. We found that host defense genes are enriched in positive selection signals, with selected variants primarily involved in regulatory functions. We also found that directional selection has operated on at least four leukocytic traits (|r| <0.6; **Figure S5**) in recent millennia. Leukocytic lineages have undergone positive selection, with the exception of eosinophils, which have decreased in proportion among granulocytes, possibly reflecting an evolutionary trade-off in favor of neutrophils, consistent with the apparently less essential role of eosinophils in immunity to infection.^71^ Conversely, non-leukocytic lineages, such as red blood cells, seem to have undergone changes in efficacy, given the observed decrease in the number of reticulocytes but the increase in their mean size and in the concentration of hemoglobin per red blood cell. These results suggest that recent positive selection has targeted the regulatory machinery underlying immune cell variation, possibly as a result of temporal changes in pathogen exposure.^35,72,73^

The estimated times of selection onset highlight the importance of the post-Neolithic period in the adaptive history of Europeans, as most selection events — both positive and negative — postdate the beginning of the Bronze Age (<4,500 ya). Furthermore, our results support a history of selection preferentially targeting variants that were already segregating in Europe before the arrival of Pontic-Caspian groups.^74^ The increase in adaptation following the Neolithic may be due to the population growth that followed the ‘Neolithic decline’,^75^ with higher selection efficacy.^76^ Alternatively, selection pressures may have increased during the Bronze Age; the expansion of urban communities, greater human mobility, animal husbandry^77^ and environmental changes^78^ may have favored the spread of epidemics, such as plague, as suggested by archeological and ancient microbial data.^13,72,73^

Our analyses also provide several lines of evidence supporting a significant contribution of antagonistic pleiotropy to the emergence of modern chronic diseases. First, selection over the last 10,000 years, particularly since the beginning of the Bronze Age, has led to a higher genetic risk of inflammatory gastrointestinal disorders^79^. Second, the main risk variants for IBD/CD are located close to key immunity genes (*IRF1, IKZF1*, *FUT2* and *SH2B3*), for which monogenic lesions confer susceptibility or resistance to infectious diseases.^27,65,67^ Third, we found that pleiotropic variants underlying infectious and inflammatory phenotypes have been primary targets of positive selection in recent millennia.

Finally, our study highlights the value of adopting an evolutionary genomics approach, not only to determine the legacy of past epidemics in human genome diversity, but also to identify candidate negatively selected variants potentially increasing infectious disease risk. Candidate negatively selected variants included the hypomorphic *LBP* D283G and TB-risk *TYK2* P1104A variants, together with *TLR3* L412F, which is associated with mild protection against autoimmune thyroid disease (OR = 0.93; *p* = 7.0 × 10^−12^)^80^ but with a modest increase in the risk of severe COVID-19 pneumonia,^81^ and *IL23R* R381Q, which increases IBD risk (OR = 1.93)^82^. Autosomal recessive or dominant TLR3 deficiency underlies viral diseases of the brain and lungs,^83,84^ whereas autosomal recessive IL23R deficiency underlies clinical disease caused by weakly virulent fungi and mycobacteria.^85^ It is tempting to speculate that *TLR3* L412F and *IL23R* R381Q confer predispositions to viral and mycobacterial/fungal infectious diseases, respectively.^86^ Detailed functional characterization of these variants in appropriate cell types, and the detection of new candidates from high-quality ancient genomes, will provide insight into the contribution of other variants to infectious disease susceptibility and severity.

In conclusion, this study shows that directional selection has targeted host defense genes over the last ten millennia of European history, particularly since the start of the Bronze Age, probably contributing to present-day disparities in susceptibility to infectious and inflammatory disorders.

### Limitations of the study

This study assumes population continuity since the Bronze Age, but fine-scale migrations have probably affected the European gene pool in modern times. Nevertheless, ancient DNA data suggest that no major population turnover has occurred over the last three millennia,^43,87–89^ indicating that our results should be largely robust to unmodeled discontinuity. Furthermore, the spatially heterogeneous nature of the aDNA dataset used here reduces the power to detect loci undergoing local adaptation. Likewise, the array used to generate ancient genomes, originally designed for demographic purposes, does not capture most rare variants, particularly those that became very rare, or even extinct, due to negative selection, providing a partial view of the evolutionary past of west Eurasians. Finally, given that environmental exposures may have differed in nature and intensity across Europe — during and beyond the studied timeframe — larger and denser sequence-based aDNA datasets are required to replicate our results and detect more subtle, region-specific selection events.

## METHODS

### Ancient DNA analysis of the 1240k capture dataset

We analyzed 2,632 aDNA genomes (**Table S7**) (i) originating from burial sites in western Eurasia (−9< longitude (°) <42.5 and 36< latitude (°) <70.1), (ii) including genotypic information for1,233,013 polymorphic sites, and (iii) retrieved from the V44.3: January 2021 release at https://reich.hms.harvard.edu/allen-ancient-dna-resource-aadr-downloadable-genotypes-present-day-and-ancient-dna-data. All individuals were treated as pseudo-haploid (i.e., hemizygotes for either the reference or the alternative allele). Using READ,^90^ we manually removed 95 samples that were either annotated as duplicated or found to correspond to first-degree relatives of at least one other individual with higher coverage in the dataset.

### Variant filtering

We removed variants for which the derived allele was present in fewer than five aDNA samples from the analysis. We also excluded variants absent from gnomAD v2.1.1 (ref^91^) and the 1,000 Genomes Project^92^ or for which the ancestral allele was not annotated in the 1,000 Genomes Project^92^. Finally, we included only variants with the ‘PASS’ flag in gnomAD v2.1.1. Other filters based on frequency were applied.

We controlled for potential artifacts due to undetected technical problems in this ‘capture’ dataset, by comparing the allele frequencies obtained with those from shotgun sequencing data. We processed 952 available published aDNA shotgun data (FASTQ files) with a published pipeline^88^ to obtain pseudohaploid data comparable with those of the capture dataset. We combined all samples and retrieved 4,620,071 variants after filtering, accounting for 95% of those present in the capture dataset. Strikingly, nine of the top 10 variants in the capture dataset, ranked by *s* value, had a frequency trajectory different from that of the shotgun dataset, with a single, strong change in frequency in the present generation, suggesting a misestimation of frequency in either the ancient or modern dataset. For these variants, we found that, when both genotypes were called in the capture and shotgun datasets, they were consistent between datasets, but there was a high percentage of missingness in the capture dataset when shotgun-sequenced individuals were called as homozygotes for the derived allele (**Table S8**). This was not the case for genotypes called as homozygous for the ancestral allele in the shotgun dataset, generating a frequency bias at population level. While searching for a pattern common to these variants, we found that only 3% of all variants in the capture dataset were within or close to an indel (within 20 bp) with a frequency >5% in European populations, 90% and 44% of our top 10 and top 100 variants from the capture dataset, ranked by *s* value, respectively, were located in such a position. We therefore conservatively removed all SNPs close to (<50 bp) an indel with a frequency >5% in the European population from the capture dataset, and variants for which the increase (or decrease) in frequency over the last 1,000 years exceeded 10% (a highly unlikely scenario), thereby excluding 28,493 variants in total. Our final dataset consisted of 933,781 SNPs from 2,537 ancient genomes.

### Allele frequency trajectories across epochs

For each SNP, we used the ancient genomes and 503 modern European genomes from the 1,000 Genomes Project (CEU,GBR,FIN,TSI and IBR populations),^92^ to compute time-series data corresponding to the trajectory of allele frequencies across various time transects. The ancient genomes were grouped by well-characterized historical periods.^35^ We considered the Neolithic (8,500-5,000 ya; *n* =729 for the capture dataset), the Bronze Age (5,000-2,500 ya; *n* = 893), the Iron Age (2,500-1,250 ya; *n* = 319) and the Middle Ages (1,250-750 ya; *n* = 435). We excluded 190 samples dated to before 8,500 ya because of the small number of individuals for such a large time period. The genetic information for the variants in ancient and modern genomes was summarized in a five-dimensional frequency vector.

### Ancestry estimation

We used factor analysis^93^ to estimate ancestry proportions at the individual level. We used a merged dataset consisting of 143,081 SNPs for 363 present-day European individuals from the 1,000 Genomes Project (IBS, TSI, GBR and FIN, available at the V42.4: March 1 2020 release) and all ancient samples (see above). Before ancestry analysis, samples were imputed with the ‘LEA’ R v3.6 package, as recommended elsewhere.^93^ We imputed individuals with high levels of genome coverage (> 795,475 SNPs covered in the 1240k dataset; *n* = 421) first, to prevent bias due to the inclusion of low-coverage samples. The remaining individuals were then imputed one-by-one and added to the imputed set of high-coverage samples. Factors are interpreted as the principal components of principal component analysis (PCA), but with temporal correction for present-day and ancient samples. We set the drift parameter so as to remove the effect of time on the *K*^th^ factor (*K* = 3 here), where *K* is the number of ancestral groups considered. As source populations, we used 41 samples annotated as Mesolithic hunter-gatherers, 25 as Anatolian farmers, and 17 as Yamnaya herders.

### Demographic model

The model used includes demographic parameters, such as divergence times, effective population sizes, migration rates and exponential growth rates of continental populations (ancestral African population, and West and East Eurasians)^35^. This model also accounts for the two major migratory movements that shaped the genetic diversity of current Europeans: the arrival of Anatolian farmers in Europe around 8,500 ya, admixing with the local Mesolithic hunter-gatherers, and that of populations of Yamnaya culture from the north of the Caucasus around 4,500 ya. We accounted for the over/under representation of a particular ancestry at selected epochs, by matching simulated ancestry proportions to the mean ancestry proportions of the observed samples, as described in ref.^35^, and by performing the analyses at the SNP level taking into account their coverage in each individual.

### Forward-in-time simulations

Computer simulations of an allele evolving under the aforementioned demographic model were performed with SLiM 3 (ref.^94^), as described elsewhere.^35^ Briefly, for each simulation, three main evolutionary parameters were randomly sampled from uniform prior distributions: the age of the mutation, the time of selection onset *T* (1,000 < *T* < 10,000 ya) and selection strength, as measured by the selection coefficient *s* (−0.05 < *s* < 0 for negative selection and 0 < *s* < 0.1 for positive selection), under an additive model (*h* = 0.5)^94^. The age of the mutation was randomly selected over the last million years of human evolution (1,000 ya < Age < 1,000,000 ya) and was defined as the point at which the mutation was introduced into the model in a randomly chosen population. Each observed ancient genome was randomly sampled from simulated diploid individuals at the generation corresponding to its calibrated, radiocarbon-based age. For each sampled individual and each polymorphic site, we randomly sampled one allele to generate pseudohaploid data, mirroring the observed pseudohaploid aDNA data used. Simulated present-day European individuals were randomly drawn from the last generation of the simulated population.

### ABC estimation

We applied the ABC approach^45^ to each of the genetic variants studied, as previously used to estimate the age (*T*_age_), strength (*s*) and the time of onset (*T*) of selection for *TYK2* P1104A (ref.^35^). Parameter estimates were obtained from 400,000 simulations (for positive or negative selection) with underlying parameters drawn from predefined uniform prior distributions. Parameters were estimated from computer simulations best fitting the observed time series data for allele frequencies. Simulated and empirical time series data were described by a vector of *K* allele frequencies over *K* epochs, used as the summary statistics to fit the observed data. For each parameter, posterior distributions, point estimates (i.e., posterior mode) and the 95% CIs were obtained from the parameter values of the 1,000 simulations with simulated summary statistics the closest to the empirical ones (‘abc’ R package, method = “Loclinear”).

### Detection of selection

We used the ABC estimates of the selection coefficient and their 95% CI (under a negative or positive selection model) to detect selection acting on specific genetic variants. The empirical threshold for rejecting neutrality (i.e., type I error estimation) was determined by simulating ~500,000 neutral alleles (*s* = 0) evolving under the same demographic model. We then estimated selection coefficients and their 95% CIs for each simulated neutral variant to obtain the distribution of *s* under neutrality (i.e., the null distribution). Simulated neutral SNPs were resampled such that the simulated and observed allele frequency spectra were identical. Finally, we determined empirical thresholds at the 1% nominal level, by calculating the 99^th^ quantile (Q_99_) of the resulting *s* distributions. Rather than using the distribution of *s* to determine these thresholds, we used the empirical distribution of the lower bound of the 95% CI of the selection coefficient (*s_l_*), as a more conservative approach to providing empirical *p* values (*p*_sel_) for each SNP. This approach, like all methods for detecting selection at the variant level, is not designed to infer the distribution of fitness effects (DFE)^95^ due to the loss of true selection signals at the high conservative thresholds used to detect robust selection candidates.

### Empirical *p* value computation

The significance threshold varied with variant frequency, as expected given that low-frequency variants are less identifiable in terms of selection than more frequent variants. We therefore excluded the variants with the lowest frequencies (the lower bound of the CI of the allele frequency <2.5%), for which estimation accuracy was poor. We normalized the data by calculating the null distribution of the lower bound of the confidence interval of the selection coefficient by frequency bin, to obtain bin-dependent significance thresholds. For the analysis of negative selection, we used the following bins: [0.025-0.05]; [0.05-0.1]; [0.1-0.2] and [0.2-0.8], whereas, for positive selection, we used the following bins: [0.025-0.2]; [0.2-0.6] and [0.6-0.8]. We identified the bin to which a variant belonged by calculating, for each variant, the CI for allele frequency estimation at each epoch, according to an approximation to the normal distribution of the 95% binomial proportion CI. We obtained the maximum for the lower bound of these CI for each SNP. Finally, if this maximum lay between 2.5% and 5%, the variant was considered to belong to the bin [0.025-0.05]. The same rationale was used for the rest of the bins. We excluded from the analysis of positive selection, any variant for which the minimum higher bound of the 95% CI of the DAF was >80%, as such variants poorly matched the simulated data. We ended up with 21,129 candidate variants for positive selection, and 27,591 for negative selection (*p*_sel_ <0.01).

Finally, as the lowest level of empirical significance depends on the number of neutral simulations in each frequency bin, we approximated the empirical null distribution with a known theoretical distribution, to improve discrimination between very small *p* values. Given the shape of the empirical null distribution, we compared the null distribution to a gamma, a beta and a lognormal distribution, for which parameters were estimated with a maximum likelihood approach (R packages ‘fitdistrplus’ and ‘EnvStats’ (v. 3.6.0)). We generated a Cullen and Frey graph (kurtosis vs. skewness) with the R package ‘fitdistrplus’ (v. 3.6.0), to distinguish between our options, and obtained *p* values for the beta distribution that best adjusted the null empirical distribution.

### Time of selection onset for positively selected loci

We evaluated the shape of the distribution of the onset of selection estimated for the top 89 positively selected variants (**Figure S3A** and **Table S2**, with mean *T* estimate of 3,327 ya), by simulating sets of 89 independent variants matching the allele frequency and the selection coefficient of the most significant variant for each positively selected LD group, and the estimated onset of selection. We investigated whether the frequency trajectories based on both ancient and modern DNA samples resulted in biased *T* estimations, due to differences in genotype calling between datasets, by re-estimating *T* values for the variant with the smallest *p*_sel_ at each of the 89 candidate positively selected loci using frequencies from aDNA only (**Figure S3B**). We thus repeated the ABC estimation for frequency trajectories, but we excluded the last epoch corresponding to current frequencies.

Finally, we assessed the contribution of adaptive admixture, by averaging the Pontic Steppe proportion of all the carriers of each of the selected alleles (for different *p*_sel_ thresholds) or that of 89,000 random alleles (**Figure S4B**). We also checked that Pontic Steppe ancestry was similar between the carriers of the variant with smallest *p*_sel_ at each of the 89 candidate loci and the simulated carriers of the 1,000 simulated variants used for each estimation of the evolutionary parameters of such variants (**Figure S4A**).

### LD grouping

We took LD into account, by using a LD map for Europeans constructed from whole-genome sequence data.^96^ This metric map displays additive “linkage disequilibrium unit” (LDU) distances, which can be used to define genetic units in which variants are in strong LD. Genomic windows were then defined as non-overlapping regions of 15 LDUs, referred to as *LD groups*. This grouping generated genomic units with a mean size of 660 kb, consistent with previous studies.^43^

### Enrichment analyses for positively selected loci

We calculated enrichment for genes (IGs or GO annotations), variants (eQTLs, ASBs, GWAS variants) and variant annotations (e.g., “missense”), with 2 × 2 contingency tables with two predefined categories (e.g. variants with *p*_sel_ <0.01 *vs*. variants with *p*_sel_ ≥ 0.01; missense variants *vs*. others), from which we calculated ORs, 95% CIs and Fisher’s exact test *p* values (for cells with counts <20) or Chi-squared *p* values, with the “oddsratio” function of the R (v 3.6.0) package “epitools”. We used independent variants to determine enrichment, by pruning variants in LD with the plink command --indep-pairwise 100 10 0.6 --maf 0.01, on our aDNA dataset, thus removing variants with *r^2^* > 0.6 in 100 kb windows, using sliding windows of 10 variants. For the *HLA* region, considered to lie between hg19 coordinates 27,298,200 and 34,036,446 of chromosome 6, we used a more conservative LD pruning method considering 1,000 kb rather than 100 kb windows (plink command --indep-pairwise 1000 100 0.6 --maf 0.01), for variants with a minor allele frequency (MAF) >1%. Where indicated, we also matched the DAF distribution of the pruned dataset to that of the studied group of variants (e.g., eQTLs or GWAS variants), using 5% frequency bins.

We calculated enrichment in IGs^97^ or GO annotations,^98^ by considering, for each LD group with >9 variants (4,134 LD groups), a binary variable, indicating whether the locus included an immunity gene or a gene with a given GO annotation, respectively. This was done to eliminate spurious enrichments due to the presence of gene clusters in a given LD group. For eQTLs analyses, we used data from a meta-analysis of whole-blood *cis*-eQTLs.^99^ We also used ENCODE data^64^ and DNase hypersensitive sites^100^ to estimate enrichment in positively selected variants for ENCODE tissues. Finally, for the study of hematopoietic traits, we retrieved GWAS data for counts or proportions of different blood cell types (36 hematopoietic traits) from UK Biobank and INTERVAL study data.^14^

### Calculations of polygenic scores

For the analysis of inflammatory and infectious traits, we calculated genetic values for each ancient individual as proposed elsewhere.^40^ Specifically, we weighted the presence/absence status of the most significant GWAS Bonferroni-significant variant (*p* < 5.0 × 10^−8^) by the GWAS-estimated effect size, for each LD group. Coverage was variable across ancient samples, and some SNPs were not present in all samples. We accounted for missing information by calculating a weighted proportion in which the estimated score was divided by the maximum possible score given the SNPs present in the sample. We then used weighted (on coverage) linear regression to investigate the association between polygenic score and ancient sample age. We included ancestry as a covariate in the model, by including the first four Factor components, and the geographic location of each sample (latitude and longitude). We compared the full model to a nested model without sample age, by performing a likelihood ratio test [R anova(nested model, full model, test = ‘Chisq’)] to obtain a *p* value. For the stratified analysis, we divided the capture dataset into mutually exclusive ancestry groups. We categorized individuals as western European hunter-gatherers, Anatolian farmers, or Pontic steppe herders if they carried over 75% of the estimated respective ancestry component (for steppe individuals, we also required the individual to be <5,000 years old). We also conducted this analysis for individuals classified as being from before or after the beginning of the Bronze Age.

For the analyses of hematological traits, we used the same method, summing alleles increasing the count or the proportion of the studied hematological trait, weighted by their effect. Genetic correlations between all 36 traits were estimated using ‘ldsc’ (ref.^101^) (**Figure S5**).

For all analyses, we checked that, despite the consideration of only one variant per LD group, none of the variants were in LD with each other. We acknowledge that the PRS obtained are proxies for the actual PRS across time because, variants that have reached fixation may not be detectable by GWAS.

### Overlap between infectious and autoimmune GWAS variants

We looked for significant overlaps between aDNA variants tagging lead GWAS SNPs for infectious and autoimmune/inflammatory diseases or traits, by retrieving summary statistics for (i) 40 GWAS of infectious diseases and (ii) 30 GWAS of inflammatory/autoimmune phenotypes (**Table S4**). We identified lead infectious or autoimmune disease-associated SNPs by retaining the variant with the highest GWAS effect size (OR) in consecutive 200 kb genomic windows. We then used aDNA variants to tag (r^2^ > 0.6) the lead GWAS-significant variants (usually absent from the aDNA array) with the plink command: plink --show-tags aDNA_variant_list --list-all --tag-r2 0.6 --tag-kb 1000. We obtained a list of aDNA variants tagging infectious and/or autoimmune disease-associated lead SNPs. We then performed 1,000 samplings of random SNPs, matched for DAF and number of LD groups, from aDNA variants tagging either infectious or autoimmune traits. For each replicate, we calculated the number of overlapping variants and found that none was higher than the observed overlap (*p* < 10^−3^). We also used a different approach to test for an enrichment in pleiotropic variants, based on the pruning of aDNA-tagged SNPs (*r^2^* = 0.6 and windows of 1 Mb) and the calculation of a classic OR for the resulting list of independent tagged SNPs. We assessed the enrichment in selection signals in the observed overlapping variants, by performing a Wilcoxon test to compare the *s* distribution of the observed overlapping variants to that of the 1,000 randomly sampled controls. Finally, the PRS and respective *p* and beta values of the regression model were obtained for the lead GWAS SNPs present in the aDNA genotyping array, either for all infectious or for all autoimmune diseases considered together. PRS and the corresponding *p* and beta values for CD, IBD, COVID A2 and COVID B2 were obtained for all GWAS-significant variants, because this information was available for such phenotypes. Of note, whereas no significant genetic correlations were found between non-overlapping infectious phenotypes (e.g. COVID_A2 and COVID B2 were considered as overlapping) nor between non-overlapping inflammatory phenotypes, formal testing for genetic correlation, as done for hematopoietic traits, was not possible since for the majority of traits we only had access to the lead GWAS-significant SNPs.

### Detection of negatively selected variants

We excluded positively selected variants from our list of candidate negatively selected variants, by calculating four haplotype-based statistics used to detect recent positive selection: iHS^102^, iHH^103^, nSL^104^ and DIND^105^. This was done for all SNPs with a DAF >0.2 in Europeans of the 1,000 Genomes Project,^92^ with *selink* (http://github.com/h-e-g/selink) and a 100 kb genomic window. We computed the 99% quantiles (Q_99_) of each of the four distributions and the proportion of SNPs with scores higher than the respective Q_99_, in 200 kb sliding windows. Only windows with >9 variants were considered. We then retrieved all the genomic windows enriched for at least one of the four selection statistics, i.e., those with proportions on the top 1% of at least one of the four distributions. LD groups overlapping, completely or partially, at least one of these windows were then removed from the analysis of negative selection.

### Site-directed mutagenesis and transient transfection for LBP

Site-directed mutagenesis was performed on the LBP-WT pCMV6 plasmid (#RC221961, OriGene) with appropriate primers (**Table S9**) and the Pfu Ultra II Fusion HS DNA (#600674, Agilent) polymerase, followed by digestion with *DpnI* (#R0176L, New England Biolabs). Plasmids were amplified in NEB-10 β competent *E. coli* (#C3019H, New England Biolabs) and purified with the HiSpeed Plasmid Maxi Kit (#12663, Qiagen). Transient transfection was carried out in HEK293T cells transfected with 1 μg of plasmid DNA in the presence of X-tremeGene9 DNA transfection reagent (#6365809001, Merck), according to the manufacturer’s instructions. Transfected cells were cultured at 37°C, under an atmosphere containing 5% CO_2_, in Dulbecco’s modified Eagle medium (DMEM) supplemented with 10% fetal bovine serum. After 48 h, supernatants and whole-cell lysates were collected for subsequent experiments.

### RNA isolation and RT-qPCR

Total RNA was extracted with the Quick-RNA MicroPrep Kit (#R1051, Zymo) according to the manufacturer’s instructions. Residual genomic DNA was removed by in-column DNase I digestion. We reverse-transcribed 1 μg of RNA with the High-Capacity RNA-to-cDNA Kit (#4387406, Thermo Fisher Scientific), and performed quantitative qPCR with PowerUp SYBR Green Master Mix (#A25742, Thermo Fisher Scientific) and the ViiA7 system (Thermo Fisher Scientific) with primers for LBP (PrimerBank ID 31652248c1 and 31652248c2) and GAPDH (PrimerBank ID 378404907c1) obtained from the PrimerBank database.^106^ Normalization of *LBP* mRNA was performed for each sample with *GAPDH* (ΔcT) and values are expressed as 2^−ΔcT^.

### Protein isolation and western blotting

Whole-cell protein lysates were extracted in modified radioimmunoprecipitation assay buffer supplemented with protease inhibitors (#5892970001, Merck) and phosphatase inhibitor cocktail (#4906837001, Merck), 0.1 mM dithiothreitol (DTT; Life Technologies), and 1 mM PMSF (#10837091001, Merck). Protein extracts and supernatants were resolved by electrophoresis in Criterion TGX 10% precast gels (Bio-Rad), with the resulting bands transferred onto PVDF membranes (#1704157, Bio-Rad) with the Transblot turbo system (Bio-Rad). Membranes were probed by incubation for 1 hour at room temperature with antibodies against LBP (#AF870-SP, R&D Systems, 1:2,000), DDK (#A8592, Merck, 1:10,000) and GAPDH (#sc-47724, Santa Cruz Biotechnology, 1:5,000). Proteins were detected by chemiluminescence with Clarity Western ECL substrate (#1705061, Bio-Rad) reagents.

### ELISA for LBP and bacterial ligand binding

Supernatants from transfected HEK293T cells were analyzed for their LBP content by Human LBP DuoSet ELISA (#DY870-05, R&D Systems) according to the manufacturer’s instructions. Bacterial ligand binding was assessed on microtiter plates coated with 30 μg/mL LPS derived from *E. coli* O111:B4 (#LPS25, Merck) or Pam2CSK4 (#tlrl-pm2s-1, InvivoGen) in 100 mM Na2CO3 (pH 9.6) for 18 hours at 4°C. The plates were blocked by incubation with 0.005% Tween and 1% bovine serum albumin in PBS for 1 hour at room temperature. Plates were then incubated with cell supernatants at various concentrations. Bound LBP was detected with the detection antibodies from the Human LBP DuoSet ELISA kit. Absorbance was read at 450 nm with a Victor X4 plate reader (Perkin Elmer).

## SUPPLEMENTAL INFORMATION

Supplemental Data include 5 figures and 9 tables and can be found with this article online at http://dx.doi.org/XXX.

## AUTHOR CONTRIBUTIONS

G.K., E.P, G.L and L.Q.M conceived and designed the study. G.K was the lead analyst, with important contributions from E.P. and G.L. A.L.N performed the functional analyses of the *LBP* D283G variant. E.P, G.L and L.Q.M oversaw the study. G.K., E.P, G.L and L.Q.M wrote the manuscript with substantial contributions from A.L.N., L.A and J.L.C.

## DECLARATION OF INTERESTS

The authors have no competing interests to declare.

## ACKNOWLEDGMENTS

We thank all members of the Human Evolutionary Genetics Laboratory at Institut Pasteur, Paris, and Nicolas Rascovan, Bertrand Boisson and Iain Mathieson for data sharing and helpful discussions. This work was supported by the *Institut Pasteur*, the *Collège de France*,the *Centre Nationale de la Recherche Scientifique* (CNRS), the *Agence Nationale de la Recherche* (ANR) grants LIFECHANGE (ANR 17 CE12 0018 02), CNSVIRGEN (ANR-19-CE15-0009-02) and MORTUI (ANR-19-CE35-0005), the French Government’s *Investissement d’Avenir* program, *Laboratoires d’Excellence* “Integrative Biology of Emerging Infectious Diseases” (ANR-10-LABX-62-IBEID) and *“Milieu Intérieur”* (ANR-10-LABX-69-01), the *Fondation pour la Recherche Médicale* (Equipe FRM DEQ20180339214), the *Fondation Allianz-Institut de France*, and the *Fondation de France* (no. 00106080). G.K. is supported by a Pasteur-Roux-Cantarini fellowship.

## DATA AND CODE AVAILABILITY

Pseudohaploid ancient and modern genome capture data are available from https://reich.hms.harvard.edu/allen-ancient-dna-resource-aadr-downloadable-genotypes-present-day-and-ancient-dna-data (V44.2). *selink* software is available from https://github.com/h-e-g/selink. The code for reproducing the figures of the paper is available from https://github.com/h-e-g/SLiM_aDNA_selection.

## WEB RESOURCES SECTION

OMIM (Online Mendelian Inheritance in Man): http://www.omim.org

eQTLS (genQTL): https://www.eqtlgen.org

GWAS atlas: https://atlas.ctglab.nl/PheWAS

Protein Atlas: https://www.proteinatlas.org/

ADASTRA: https://adastra.autosome.ru

Open Targets Genetics: https://genetics.opentargets.org

GWAS catalog: https://www.ebi.ac.uk/gwas/home

## Notes

### Competing Interest Statement

The authors have declared no competing interest.

## REFERENCES

1. Casanova, J.-L., and Abel, L. (2005). Inborn errors of immunity to infection : the rule rather than the exception. J Exp Med 202, 197–201. 10.1084/jem.20050854.

2. Cairns, J. (1997). Matters of Life and Death (Princeton University Press).

3. Allison, A.C. (1954). Protection afforded by sickle-cell trait against subtertian malareal infection. Br Med J 1, 290–294. 10.1136/bmj.1.4857.290.

4. Barreiro, L.B., and Quintana-Murci, L. (2010). From evolutionary genetics to human immunology: how selection shapes host defence genes. Nat Rev Genet 11, 17–30. 10.1038/nrg2698.

5. Quintana-Murci, L. (2019). Human Immunology through the Lens of Evolutionary Genetics. Cell 177, 184–199. https://doi.org/10.1016/j.cell.2019.02.033.

6. Fumagalli, M., and Sironi, M. (2014). Human genome variability, natural selection and infectious diseases. Curr Opin Immunol 30C, 9–16.

7. Karlsson, E.K., Kwiatkowski, D.P., and Sabeti, P.C. (2014). Natural selection and infectious disease in human populations. Nat Rev Genet 15, 379–393. 10.1038/nrg3734.

8. Quintana-Murci, L., and Clark, A.G. (2013). Population genetic tools for dissecting innate immunity in humans. Nat Rev Immunol 13, 280–293. 10.1038/nri3421.

9. Diamond, J. (2002). Evolution, consequences and future of plant and animal domestication. Nature 418, 700–707. 10.1038/nature01019.

10. Key, F.M., Posth, C., Esquivel-Gomez, L.R., Hübler, R., Spyrou, M.A., Neumann, G.U., Furtwängler, A., Sabin, S., Burri, M., Wissgott, A., et al. (2020). Emergence of human-adapted Salmonella enterica is linked to the Neolithization process. Nat Ecol Evol 4, 324–333. 10.1038/s41559-020-1106-9.

11. Wolfe, N.D., Dunavan, C.P., and Diamond, J. (2007). Origins of major human infectious diseases. Nature 447, 279–283. 10.1038/nature05775.

12. Harper, K.N., and Armelagos, G.J. (2013). Genomics, the origins of agriculture, and our changing microbe-scape: time to revisit some old tales and tell some new ones. Am J Phys Anthropol 152 Suppl 57, 135–152. 10.1002/ajpa.22396.

13. Fuchs, K., Rinne, C., Drummer, C., Immel, A., Krause-Kyora, B., and Nebel, A. (2019). Infectious diseases and Neolithic transformations: Evaluating biological and archaeological proxies in the German loess zone between 5500 and 2500 BCE. The Holocene 29, 1545–1557. 10.1177/0959683619857230.

14. Astle, W.J., Elding, H., Jiang, T., Allen, D., Ruklisa, D., Mann, A.L., Mead, D., Bouman, H., Riveros-Mckay, F., Kostadima, M.A., et al. (2016). The Allelic Landscape of Human Blood Cell Trait Variation and Links to Common Complex Disease. Cell 167, 1415–1429.e1419. 10.1016/j.cell.2016.10.042.

15. Bao, E.L., Cheng, A.N., and Sankaran, V.G. (2019). The genetics of human hematopoiesis and its disruption in disease. EMBO Mol Med 11, e10316–e10316. 10.15252/emmm.201910316.

16. Liggett, L.A., and Sankaran, V.G. (2020). Unraveling Hematopoiesis through the Lens of Genomics. Cell 182, 1384–1400. 10.1016/j.cell.2020.08.030.

17. Notarangelo, L.D., Bacchetta, R., Casanova, J.-L., and Su, H.C. (2020). Human inborn errors of immunity: An expanding universe. Sci Immunol 5, eabb1662. 10.1126/sciimmunol.abb1662.

18. Riley, J.C. (2001). Rising Life Expectancy: A Global History (Cambridge University Press). DOI: 10.1017/CBO9781316036495.

19. Barreiro, L.B., and Quintana-Murci, L. (2020). Evolutionary and population (epi)genetics of immunity to infection. Hum Genet 139, 723–732. 10.1007/s00439-020-02167-x.

20. Benton, M.L., Abraham, A., LaBella, A.L., Abbot, P., Rokas, A., and Capra, J.A. (2021). The influence of evolutionary history on human health and disease. Nat Rev Genet 22, 269–283. 10.1038/s41576-020-00305-9.

21. Sironi, M., and Clerici, M. (2010). The hygiene hypothesis: an evolutionary perspective. Microbes and Infection 12, 421–427. https://doi.org/10.1016/j.micinf.2010.02.002.

22. Fodil, N., Langlais, D., and Gros, P. (2016). Primary Immunodeficiencies and Inflammatory Disease: A Growing Genetic Intersection. Trends Immunol 37, 126–140. 10.1016/j.it.2015.12.006.

23. Langlais, D., Fodil, N., and Gros, P. (2017). Genetics of Infectious and Inflammatory Diseases: Overlapping Discoveries from Association and Exome-Sequencing Studies. Ann Rev Immunol 35, 1–30. 10.1146/annurev-immunol-051116-052442.

24. Fumagalli, M., Pozzoli, U., Cagliani, R., Comi, G.P., Riva, S., Clerici, M., Bresolin, N., and Sironi, M. (2009). Parasites represent a major selective force for interleukin genes and shape the genetic predisposition to autoimmune conditions. J Exp Med 206, 1395–1408. 10.1084/jem.20082779.

25. Jostins, L., Ripke, S., Weersma, R.K., Duerr, R.H., McGovern, D.P., Hui, K.Y., Lee, J.C., Schumm, L.P., Sharma, Y., Anderson, C.A., et al. (2012). Host-microbe interactions have shaped the genetic architecture of inflammatory bowel disease. Nature 491, 119–124. 10.1038/nature11582.

26. Tsoi, L.C., Spain, S.L., Knight, J., Ellinghaus, E., Stuart, P.E., Capon, F., Ding, J., Li, Y., Tejasvi, T., Gudjonsson, J.E., et al. (2012). Identification of 15 new psoriasis susceptibility loci highlights the role of innate immunity. Nat Genet 44, 1341–1348. 10.1038/ng.2467.

27. Zhernakova, A., Elbers, C.C., Ferwerda, B., Romanos, J., Trynka, G., Dubois, P.C., de Kovel, C.G.F., Franke, L., Oosting, M., Barisani, D., et al. (2010). Evolutionary and functional analysis of celiac risk loci reveals SH2B3 as a protective factor against bacterial infection. Am J Hum Genet 86, 970–977. 10.1016/j.ajhg.2010.05.004.

28. Prugnolle, F., Manica, A., Charpentier, M., Guegan, J.F., Guernier, V., and Balloux, F. (2005). Pathogen-driven selection and worldwide HLA class I diversity. Curr Biol 15, 1022–1027. 10.1016/j.cub.2005.04.050.

29. Chen, H., Hayashi, G., Lai, O.Y., Dilthey, A., Kuebler, P.J., Wong, T.V., Martin, M.P., Fernandez Vina, M.A., McVean, G., Wabl, M., et al. (2012). Psoriasis patients are enriched for genetic variants that protect against HIV-1 disease. PLoS Genet 8, e1002514–e1002514. 10.1371/journal.pgen.1002514.

30. Gough, S.C.L., and Simmonds, M.J. (2007). The HLA Region and Autoimmune Disease: Associations and Mechanisms of Action. Curr Genomics 8, 453–465. 10.2174/138920207783591690.

31. Matzaraki, V., Kumar, V., Wijmenga, C., and Zhernakova, A. (2017). The MHC locus and genetic susceptibility to autoimmune and infectious diseases. Genome Biol 18, 76. 10.1186/s13059-017-1207-1.

32. Ritari, J., Koskela, S., Hyvärinen, K., FinnGen, and Partanen, J. (2022). HLA-disease association and pleiotropy landscape in over 235,000 Finns. Hum Immunol 83, 391–398. https://doi.org/10.1016/j.humimm.2022.02.003.

33. Ryder, L.P., Svejgaard, A., and Dausset, J. (1981). GENETICS OF HLA DISEASE ASSOCIATION. Ann Rev Genet 15, 169–187. 10.1146/annurev.ge.15.120181.001125.

34. Boisson-Dupuis, S., Ramirez-Alejo, N., Li, Z., Patin, E., Rao, G., Kerner, G., Lim, C.K., Krementsov, D.N., Hernandez, N., Ma, C.S., et al. (2018). Tuberculosis and impaired IL-23-dependent IFN-γ immunity in humans homozygous for a common TYK2 missense variant. Sci Immunol 3, eaau8714. 10.1126/sciimmunol.aau8714.

35. Kerner, G., Laval, G., Patin, E., Boisson-Dupuis, S., Abel, L., Casanova, J.-L., and Quintana-Murci, L. (2021). Human ancient DNA analyses reveal the high burden of tuberculosis in Europeans over the last 2,000 years. Am J Hum Genet 108, 517–524. 10.1016/j.ajhg.2021.02.009.

36. Dendrou, C.A., Cortes, A., Shipman, L., Evans, H.G., Attfield, K.E., Jostins, L., Barber, T., Kaur, G., Kuttikkatte, S.B., Leach, O.A., et al. (2016). Resolving TYK2 locus genotype-to-phenotype differences in autoimmunity. Sci Transl Med 8, 363ra149–363ra149. 10.1126/scitranslmed.aag1974.

37. Diogo, D., Bastarache, L., Liao, K.P., Graham, R.R., Fulton, R.S., Greenberg, J.D., Eyre, S., Bowes, J., Cui, J., Lee, A., et al. (2015). TYK2 protein-coding variants protect against rheumatoid arthritis and autoimmunity, with no evidence of major pleiotropic effects on non-autoimmune complex traits. PLoS One 10, e0122271–e0122271. 10.1371/journal.pone.0122271.

38. International Genetics of Ankylosing Spondylitis, C., Cortes, A., Hadler, J., Pointon, J.P., Robinson, P.C., Karaderi, T., Leo, P., Cremin, K., Pryce, K., Harris, J., et al. (2013). Identification of multiple risk variants for ankylosing spondylitis through high-density genotyping of immune-related loci. Nat Genet 45, 730–738. 10.1038/ng.2667.

39. Kerner, G., Ramirez-Alejo, N., Seeleuthner, Y., Yang, R., Ogishi, M., Cobat, A., Patin, E., Quintana-Murci, L., Boisson-Dupuis, S., Casanova, J.-L., and Abel, L. (2019). Homozygosity for TYK2 P1104A underlies tuberculosis in about 1% of patients in a cohort of European ancestry. Proc Natl Acad Sci U S A 116, 10430–10434. 10.1073/pnas.1903561116.

40. Ju, D., and Mathieson, I. (2021). The evolution of skin pigmentation-associated variation in West Eurasia. Proc Natl Acad Sci U S A 118, e2009227118. 10.1073/pnas.2009227118.

41. Key, F.M., Fu, Q., Romagné, F., Lachmann, M., and Andrés, A.M. (2016). Human adaptation and population differentiation in the light of ancient genomes. Nat Commun 7, 10775–10775. 10.1038/ncomms10775.

42. Mathieson, I. (2020). Limited Evidence for Selection at the FADS Locus in Native American Populations. Mol Biol Evol 37, 2029–2033. 10.1093/molbev/msaa064.

43. Mathieson, I., Lazaridis, I., Rohland, N., Mallick, S., Patterson, N., Roodenberg, S.A., Harney, E., Stewardson, K., Fernandes, D., Novak, M., et al. (2015). Genome-wide patterns of selection in 230 ancient Eurasians. Nature 528, 499–503. 10.1038/nature16152.

44. Lindo, J., Huerta-Sánchez, E., Nakagome, S., Rasmussen, M., Petzelt, B., Mitchell, J., Cybulski, J.S., Willerslev, E., DeGiorgio, M., and Malhi, R.S. (2016). A time transect of exomes from a Native American population before and after European contact. Nat Commun 7, 13175–13175. 10.1038/ncomms13175.

45. Beaumont, M.A., and Rannala, B. (2004). The Bayesian revolution in genetics. Nat Rev Genet 5, 251–261. 10.1038/nrg1318.

46. Lazaridis, I., Patterson, N., Mittnik, A., Renaud, G., Mallick, S., Kirsanow, K., Sudmant, P.H., Schraiber, J.G., Castellano, S., Lipson, M., et al. (2014). Ancient human genomes suggest three ancestral populations for present-day Europeans. Nature 513, 409–413. 10.1038/nature13673.

47. Skoglund, P., Malmström, H., Raghavan, M., Storå, J., Hall, P., Willerslev, E., Gilbert, M.T.P., Götherström, A., and Jakobsson, M. (2012). Origins and Genetic Legacy of Neolithic Farmers and Hunter-Gatherers in Europe. Science 336, 466–469. 10.1126/science.1216304.

48. Allentoft, M.E., Sikora, M., Sjögren, K.-G., Rasmussen, S., Rasmussen, M., Stenderup, J., Damgaard, P.B., Schroeder, H., Ahlström, T., Vinner, L., et al. (2015). Population genomics of Bronze Age Eurasia. Nature 522, 167–172. 10.1038/nature14507.

49. Haak, W., Lazaridis, I., Patterson, N., Rohland, N., Mallick, S., Llamas, B., Brandt, G., Nordenfelt, S., Harney, E., Stewardson, K., et al. (2015). Massive migration from the steppe was a source for Indo-European languages in Europe. Nature 522, 207–211. 10.1038/nature14317.

50. Burger, J., Link, V., Blöcher, J., Schulz, A., Sell, C., Pochon, Z., Diekmann, Y., Žegarac, A., Hofmanová, Z., Winkelbach, L., et al. (2020). Low Prevalence of Lactase Persistence in Bronze Age Europe Indicates Ongoing Strong Selection over the Last 3,000 Years. Curr Biol 30, 4307–4315.e4313. https://doi.org/10.1016/j.cub.2020.08.033.

51. Segurel, L., Guarino-Vignon, P., Marchi, N., Lafosse, S., Laurent, R., Bon, C., Fabre, A., Hegay, T., and Heyer, E. (2020). Why and when was lactase persistence selected for? Insights from Central Asian herders and ancient DNA. PLoS Biol 18, e3000742–e3000742. 10.1371/journal.pbio.3000742.

52. Kristiansen, H., Scherer, C.A., McVean, M., Iadonato, S.P., Vends, S., Thavachelvam, K., Steffensen, T.B., Horan, K.A., Kuri, T., Weber, F., et al. (2010). Extracellular 2’-5’ oligoadenylate synthetase stimulates RNase L-independent antiviral activity: a novel mechanism of virus-induced innate immunity. J Virol 84, 11898–11904. 10.1128/JVI.01003-10.

53. Shelton, J.F., Shastri, A.J., Ye, C., Weldon, C.H., Filshtein-Sonmez, T., Coker, D., Symons, A., Esparza-Gordillo, J., Chubb, A., Fitch, A., et al. (2021). Trans-ancestry analysis reveals genetic and nongenetic associations with COVID-19 susceptibility and severity. Nat Genet 53, 801–808. 10.1038/s41588-021-00854-7.

54. Pickrell, J.K., Berisa, T., Liu, J.Z., Ségurel, L., Tung, J.Y., and Hinds, D.A. (2016). Detection and interpretation of shared genetic influences on 42 human traits. Nat Genet 48, 709–717. 10.1038/ng.3570.

55. Tian, C., Hromatka, B.S., Kiefer, A.K., Eriksson, N., Noble, S.M., Tung, J.Y., and Hinds, D.A. (2017). Genome-wide association and HLA region fine-mapping studies identify susceptibility loci for multiple common infections. Nat Commun 8, 599–599. 10.1038/s41467-017-00257-5.

56. Malaria Genomic Epidemiology, N. (2019). Insights into malaria susceptibility using genome-wide data on 17,000 individuals from Africa, Asia and Oceania. Nat Commun 10, 5732–5732. 10.1038/s41467-019-13480-z.

57. Ségurel, L., Thompson, E.E., Flutre, T., Lovstad, J., Venkat, A., Margulis, S.W., Moyse, J., Ross, S., Gamble, K., Sella, G., et al. (2012). The ABO blood group is a trans-species polymorphism in primates. Proc Natl Acad Sci U S A 109, 18493–18498. 10.1073/pnas.1210603109.

58. Cuadros-Espinoza, S., Laval, G., Quintana-Murci, L., and Patin, E. (2022). The genomic signatures of natural selection in admixed human populations. Am J Hum Genet 109, 710–726. https://doi.org/10.1016/j.ajhg.2022.02.011.

59. Zeberg, H., and Pääbo, S. (2021). A genomic region associated with protection against severe COVID-19 is inherited from Neandertals. Proc Natl Acad Sci U S A 118, e2026309118. doi:10.1073/pnas.2026309118.

60. Huffman, J.E., Butler-Laporte, G., Khan, A., Pairo-Castineira, E., Drivas, T.G., Peloso, G.M., Nakanishi, T., Initiative, C.-H.G., Ganna, A., Verma, A., et al. (2022). Multi-ancestry fine mapping implicates OAS1 splicing in risk of severe COVID-19. Nat Genet 54, 125–127. 10.1038/s41588-021-00996-8.

61. Zhou, S., Butler-Laporte, G., Nakanishi, T., Morrison, D.R., Afilalo, J., Afilalo, M., Laurent, L., Pietzner, M., Kerrison, N., Zhao, K., et al. (2021). A Neanderthal OAS1 isoform protects individuals of European ancestry against COVID-19 susceptibility and severity. Nat Med 27, 659–667. 10.1038/s41591-021-01281-1.

62. Abramov, S., Boytsov, A., Bykova, D., Penzar, D.D., Yevshin, I., Kolmykov, S.K., Fridman, M.V., Favorov, A.V., Vorontsov, I.E., Baulin, E., et al. (2021). Landscape of allele-specific transcription factor binding in the human genome. Nat Commun 12, 2751–2751. 10.1038/s41467-021-23007-0.

63. Ferreira, M.A.R., Mathur, R., Vonk, J.M., Szwajda, A., Brumpton, B., Granell, R., Brew, B.K., Ullemar, V., Lu, Y., Jiang, Y., et al. (2019). Genetic Architectures of Childhood- and Adult-Onset Asthma Are Partly Distinct. Am J Hum Genet 104, 665–684. 10.1016/j.ajhg.2019.02.022.

64. Davis, C.A., Hitz, B.C., Sloan, C.A., Chan, E.T., Davidson, J.M., Gabdank, I., Hilton, J.A., Jain, K., Baymuradov, U.K., Narayanan, A.K., et al. (2018). The Encyclopedia of DNA elements (ENCODE): data portal update. Nucleic Acids Res 46, D794–D801. 10.1093/nar/gkx1081.

65. Nunes-Santos, C.J., Kuehn, H.S., and Rosenzweig, S.D. (2020). IKAROS Family Zinc Finger 1-Associated Diseases in Primary Immunodeficiency Patients. Immunol Allergy Clin North Am 40, 461–470. 10.1016/j.iac.2020.04.004.

66. Panda, D., Gjinaj, E., Bachu, M., Squire, E., Novatt, H., Ozato, K., and Rabin, R.L. (2019). IRF1 Maintains Optimal Constitutive Expression of Antiviral Genes and Regulates the Early Antiviral Response. Front Immunol 10. 10.3389/fimmu.2019.01019.

67. Lindesmith, L., Moe, C., Marionneau, S., Ruvoen, N., Jiang, X., Lindblad, L., Stewart, P., LePendu, J., and Baric, R. (2003). Human susceptibility and resistance to Norwalk virus infection. Nat Med 9, 548–553. 10.1038/nm860.

68. Raza, M.W., Blackwell, C.C., Molyneaux, P., James, V.S., Ogilvie, M.M., Inglis, J.M., and Weir, D.M. (1991). Association between secretor status and respiratory viral illness. BMJ 303, 815–818. 10.1136/bmj.303.6806.815.

69. Park, B.S., and Lee, J.-O. (2013). Recognition of lipopolysaccharide pattern by TLR4 complexes. Exp Mol Med 45, e66–e66. 10.1038/emm.2013.97.

70. Eckert, Jana K., Kim, Young J., Kim, Jung I., Gürtler, K., Oh, D.-Y., Sur, S., Lundvall, L., Hamann, L., van der Ploeg, A., Pickkers, P., et al. (2013). The Crystal Structure of Lipopolysaccharide Binding Protein Reveals the Location of a Frequent Mutation that Impairs Innate Immunity. Immunity 39, 647–660. https://doi.org/10.1016/j.immuni.2013.09.005.

71. Gleich, G.J., Klion, A.D., Lee, J.J., and Weller, P.F. (2013). The consequences of not having eosinophils. Allergy 68, 829–835. 10.1111/all.12169.

72. Andrades Valtueña, A., Neumann Gunnar, U., Spyrou Maria, A., Musralina, L., Aron, F., Beisenov, A., Belinskiy Andrey, B., Bos Kirsten, I., Buzhilova, A., Conrad, M., et al. (2022). Stone Age Yersinia pestis genomes shed light on the early evolution, diversity, and ecology of plague. Proc Natl Acad Sci U S A 119, e2116722119. 10.1073/pnas.2116722119.

73. Rascovan, N., Sjögren, K.-G., Kristiansen, K., Nielsen, R., Willerslev, E., Desnues, C., and Rasmussen, S. (2019). Emergence and Spread of Basal Lineages of Yersinia pestis during the Neolithic Decline. Cell 176, 295–305.e210. https://doi.org/10.1016/j.cell.2018.11.005.

74. Skoglund, P., and Mathieson, I. (2018). Ancient Genomics of Modern Humans: The First Decade. Ann Rev Genom Hum Genet 19, 381–404. 10.1146/annurev-genom-083117-021749.

75. Kristiansen, K., Fowler, C., Harding, J., & Hofmann, D. (2014). The Decline of the Neolithic and the Rise of Bronze Age Society. Oxford Handbooks Online. https://doi.org/10.1093/OXFORDHB/9780199545841.013.057.

76. Charlesworth, B. (2009). Effective population size and patterns of molecular evolution and variation. Nat Rev Genet 10, 195–205. 10.1038/nrg2526.

77. Scott, A., Reinhold, S., Hermes, T., Kalmykov, A.A., Belinskiy, A., Buzhilova, A., Berezina, N., Kantorovich, A.R., Maslov, V.E., Guliyev, F., et al. (2022). Emergence and intensification of dairying in the Caucasus and Eurasian steppes. Nat Ecol Evol. 10.1038/s41559-022-01701-6.

78. Racimo, F., Woodbridge, J., Fyfe, R.M., Sikora, M., Sjögren, K.-G., Kristiansen, K., and Vander Linden, M. (2020). The spatiotemporal spread of human migrations during the European Holocene. Proc Natl Acad Sci U S A 117, 8989–9000. 10.1073/pnas.1920051117.

79. Song, W., Shi, Y., Wang, W., Pan, W., Qian, W., Yu, S., Zhao, M., and Lin, G.N. (2021). A selection pressure landscape for 870 human polygenic traits. Nat Hum Behav 5, 1731–1743. 10.1038/s41562-021-01231-4.

80. Saevarsdottir, S., Olafsdottir, T.A., Ivarsdottir, E.V., Halldorsson, G.H., Gunnarsdottir, K., Sigurdsson, A., Johannesson, A., Sigurdsson, J.K., Juliusdottir, T., Lund, S.H., et al. (2020). FLT3 stop mutation increases FLT3 ligand level and risk of autoimmune thyroid disease. Nature 584, 619–623. 10.1038/s41586-020-2436-0.

81. Croci, S., Venneri, M.A., Mantovani, S., Fallerini, C., Benetti, E., Picchiotti, N., Campolo, F., Imperatore, F., Palmieri, M., Daga, S., et al. (2021). The polymorphism L412F in TLR3 inhibits autophagy and is a marker of severe COVID-19 in males. Autophagy, 1–11. 10.1080/15548627.2021.1995152.

82. de Lange, K.M., Moutsianas, L., Lee, J.C., Lamb, C.A., Luo, Y., Kennedy, N.A., Jostins, L., Rice, D.L., Gutierrez-Achury, J., Ji, S.-G., et al. (2017). Genome-wide association study implicates immune activation of multiple integrin genes in inflammatory bowel disease. Nat Genet 49, 256–261. 10.1038/ng.3760.

83. Guo, Y., Audry, M., Ciancanelli, M., Alsina, L., Azevedo, J., Herman, M., Anguiano, E., Sancho-Shimizu, V., Lorenzo, L., Pauwels, E., et al. (2011). Herpes simplex virus encephalitis in a patient with complete TLR3 deficiency: TLR3 is otherwise redundant in protective immunity. J Exp Med 208, 2083–2098. 10.1084/jem.20101568.

84. Zhang, S.Y., Jouanguy, E., Ugolini, S., Smahi, A., Elain, G., Romero, P., Segal, D., Sancho-Shimizu, V., Lorenzo, L., Puel, A., et al. (2007). TLR3 deficiency in patients with herpes simplex encephalitis. Science 317, 1522–1527. 317/5844/1522 [pii]10.1126/science. 1139522.

85. Martínez-Barricarte, R., Markle, J.G., Ma, C.S., Deenick, E.K., Ramírez-Alejo, N., Mele, F., Latorre, D., Mahdaviani, S.A., Aytekin, C., Mansouri, D., et al. (2018). Human IFN-γ immunity to mycobacteria is governed by both IL-12 and IL-23. Sci Immunol 3, eaau6759. 10.1126/sciimmunol.aau6759.

86. Sun, R., Hedl, M., and Abraham, C. (2020). IL23 induces IL23R recycling and amplifies innate receptor-induced signalling and cytokines in human macrophages, and the IBD-protective IL23R R381Q variant modulates these outcomes. Gut 69, 264–273. 10.1136/gutjnl-2018-316830.

87. Brunel, S., Bennett, E.A., Cardin, L., Garraud, D., Emam, H.B., Beylier, A., Boulestin, B., Chenal, F., Ciesielski, E., Convertini, F., et al. (2020). Ancient genomes from present-day France unveil 7,000 years of its demographic history. Proc Natl Acad Sci U S A 117, 12791–12798. doi:10.1073/pnas.1918034117.

88. Margaryan, A., Lawson, D.J., Sikora, M., Racimo, F., Rasmussen, S., Moltke, I., Cassidy, L.M., Jørsboe, E., Ingason, A., Pedersen, M.W., et al. (2020). Population genomics of the Viking world. Nature 585, 390–396. 10.1038/s41586-020-2688-8.

89. Olalde, I., Mallick, S., Patterson, N., Rohland, N., Villalba-Mouco, V., Silva, M., Dulias, K., Edwards, C.J., Gandini, F., Pala, M., et al. (2019). The genomic history of the Iberian Peninsula over the past 8000 years. Science 363, 1230–1234. doi:10.1126/science.aav4040.

90. Monroy Kuhn, J.M., Jakobsson, M., and Günther, T. (2018). Estimating genetic kin relationships in prehistoric populations. PLoS One 13, e0195491–e0195491. 10.1371/journal.pone.0195491.

91. Karczewski, K.J., Francioli, L.C., Tiao, G., Cummings, B.B., Alfoldi, J., Wang, Q., Collins, R.L., Laricchia, K.M., Ganna, A., Birnbaum, D.P., et al. (2020). The mutational constraint spectrum quantified from variation in 141,456 humans. Nature 581, 434–443. 10.1038/s41586-020-2308-7.

92. Fairley, S., Lowy-Gallego, E., Perry, E., and Flicek, P. (2020). The International Genome Sample Resource (IGSR) collection of open human genomic variation resources. Nucleic Acids Res 48, D941–D947. 10.1093/nar/gkz836.

93. François, O., and Jay, F. (2020). Factor analysis of ancient population genomic samples. Nat Commun 11, 4661–4661. 10.1038/s41467-020-18335-6.

94. Haller, B.C., and Messer, P.W. (2019). SLiM 3: Forward Genetic Simulations Beyond the Wright-Fisher Model. Mol Biol Evol 36, 632–637. 10.1093/molbev/msy228.

95. Gutenkunst, R.N., Hernandez, R.D., Williamson, S.H., and Bustamante, C.D. (2009). Inferring the joint demographic history of multiple populations from multidimensional SNP frequency data. PLoS Genet 5, e1000695–e1000695. 10.1371/journal.pgen.1000695.

96. Vergara-Lope, A., Jabalameli, M.R., Horscroft, C., Ennis, S., Collins, A., and Pengelly, R.J. (2019). Linkage disequilibrium maps for European and African populations constructed from whole genome sequence data. Sci Data 6, 208–208. 10.1038/s41597-019-0227-y.

97. Deschamps, M., Laval, G., Fagny, M., Itan, Y., Abel, L., Casanova, J.-L., Patin, E., and Quintana-Murci, L. (2016). Genomic Signatures of Selective Pressures and Introgression from Archaic Hominins at Human Innate Immunity Genes. Am J Hum Genet 98, 5–21. 10.1016/j.ajhg.2015.11.014.

98. Young, M.D., Wakefield, M.J., Smyth, G.K., and Oshlack, A. (2010). Gene ontology analysis for RNA-seq: accounting for selection bias. Genome Biol 11, R14. 10.1186/gb-2010-11-2-r14.

99. Võsa, U., Claringbould, A., Westra, H.-J., Bonder, M.J., Deelen, P., Zeng, B., Kirsten, H., Saha, A., Kreuzhuber, R., Yazar, S., et al. (2021). Large-scale cis- and trans-eQTL analyses identify thousands of genetic loci and polygenic scores that regulate blood gene expression. Nat Genet 53, 1300–1310. 10.1038/s41588-021-00913-z.

100. Schmidt, E.M., Zhang, J., Zhou, W., Chen, J., Mohlke, K.L., Chen, Y.E., and Willer, C.J. (2015). GREGOR: evaluating global enrichment of trait-associated variants in epigenomic features using a systematic, data-driven approach. Bioinformatics 31, 2601–2606. 10.1093/bioinformatics/btv201.

101. Bulik-Sullivan, B., Finucane, H.K., Anttila, V., Gusev, A., Day, F.R., Loh, P.R., ReproGen, C., Psychiatric Genomics, C., Genetic Consortium for Anorexia Nervosa of the Wellcome Trust Case Control, C., Duncan, L., et al. (2015). An atlas of genetic correlations across human diseases and traits. Nat Genet 47, 1236–1241. 10.1038/ng.3406.

102. Voight, B.F., Kudaravalli, S., Wen, X., and Pritchard, J.K. (2006). A map of recent positive selection in the human genome. PLoS Biol 4, e72–e72. 10.1371/journal.pbio.0040072.

103. Grossman Sharon, R., Shylakhter, I., Karlsson Elinor, K., Byrne Elizabeth, H., Morales, S., Frieden, G., Hostetter, E., Angelino, E., Garber, M., Zuk, O., et al. (2010). A Composite of Multiple Signals Distinguishes Causal Variants in Regions of Positive Selection. Science 327, 883–886. 10.1126/science.1183863.

104. Ferrer-Admetlla, A., Liang, M., Korneliussen, T., and Nielsen, R. (2014). On detecting incomplete soft or hard selective sweeps using haplotype structure. Mol Biol Evol 31, 1275–1291. 10.1093/molbev/msu077.

105. Barreiro, L.B., Ben-Ali, M., Quach, H., Laval, G., Patin, E., Pickrell, J.K., Bouchier, C., Tichit, M., Neyrolles, O., Gicquel, B., et al. (2009). Evolutionary dynamics of human Toll-like receptors and their different contributions to host defense. PLoS Genet 5, e1000562–e1000562. 10.1371/journal.pgen.1000562.

106. Wang, X., Spandidos, A., Wang, H., and Seed, B. (2012). PrimerBank: a PCR primer database for quantitative gene expression analysis, 2012 update. Nucleic Acids Res 40, D1144–1149. 10.1093/nar/gkr1013.

