## Supplemental Information for "Genetic adaptation to pathogens and increased risk of inflammatory disorders in post-Neolithic Europe"

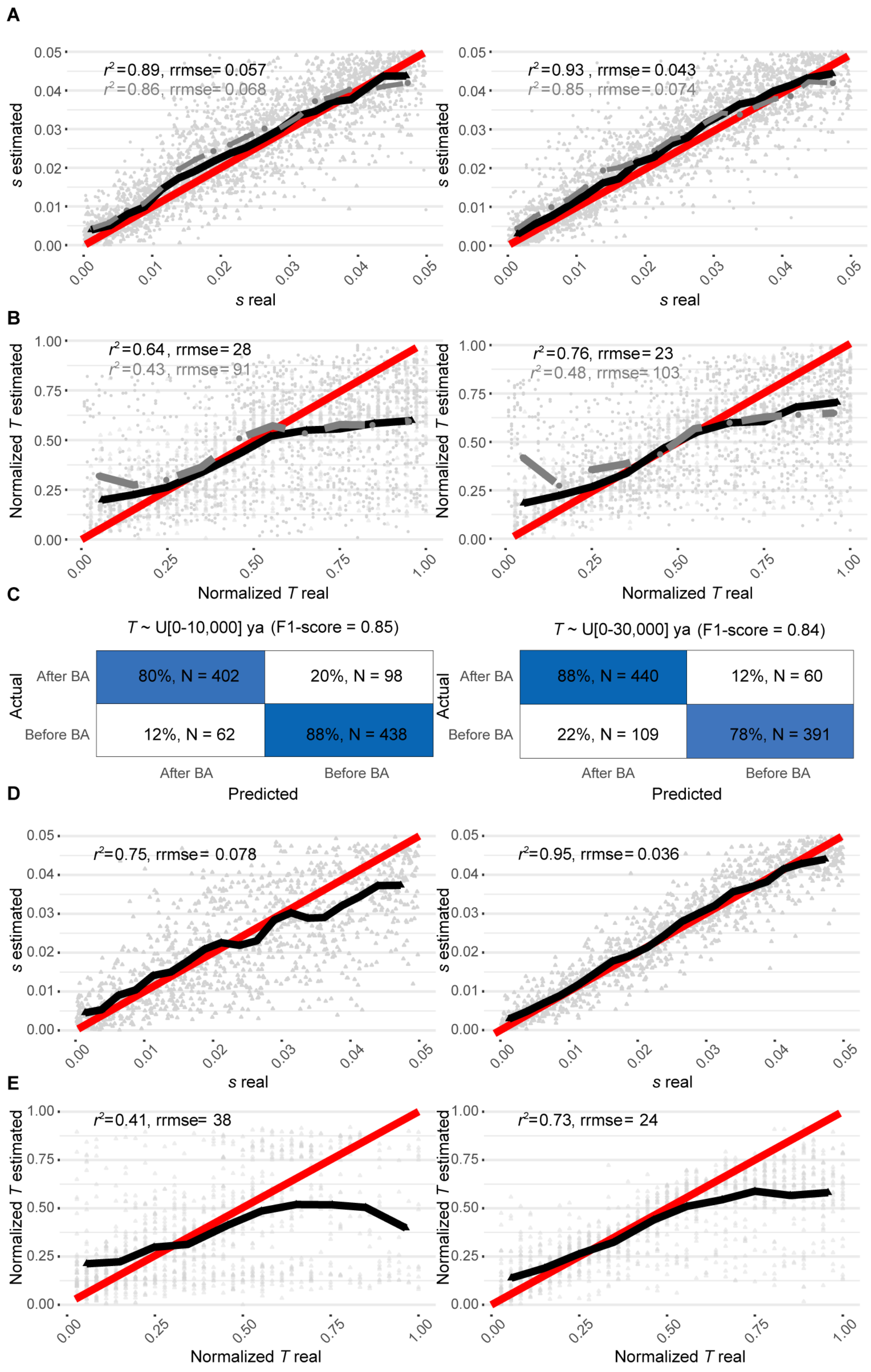


**Figure S1. Cross-validation for the time of selection onset and selection intensity**
Leave-one-out cross-validation for (A,D) the selection coefficient (*s*) or (B,E) the onset of selection (*T*), for either (A,B) positive or (D,E) negative selection. The left column shows analyses on variants for which the maximum of the lower bounds of the CI for the frequency of the variant across epochs was between 0 and 20% (**STAR Methods**) and the right column shows analyses on variants for which this maximum was between 20 and 60%. Each panel was built from $N$ = 1,000 repetitions. Red lines represent identity functions. Black continuous

lines (for simulations drawn with $T \sim$ U[-10,000-0]) with black triangles and gray dashed lines ($T \sim$ U[-30,000-0]) with gray circles indicate means for the corresponding estimated values ($y$-axis) over small intervals for the real values ($x$-axis). Gray triangles (for simulations drawn with $T \sim$ U[-10,000-0]) and gray circles ($T \sim$ U[-30,000-0]) in the background represent all1,000 values obtained for each cross-validation model. (C) Confusion matrix for 1,000 randomly chosen simulated variants drawn as in the right panel of (B) with a prior distribution of $T \sim$ U[-10,000-0], 500 with an onset of selection <4,500 ya and 500 with an onset of selection >4,500 ya (left panel), or for 1,000 randomly chosen simulated variants drawn as in the right panel of (B) with a prior distribution of $T \sim$ U[-30,000-0], 500 with an onset of selection <4,500 ya and 500 with an onset of selection >4,500 ya (right panel). The goodness-of-fit for the method is summarized as an F1-score.

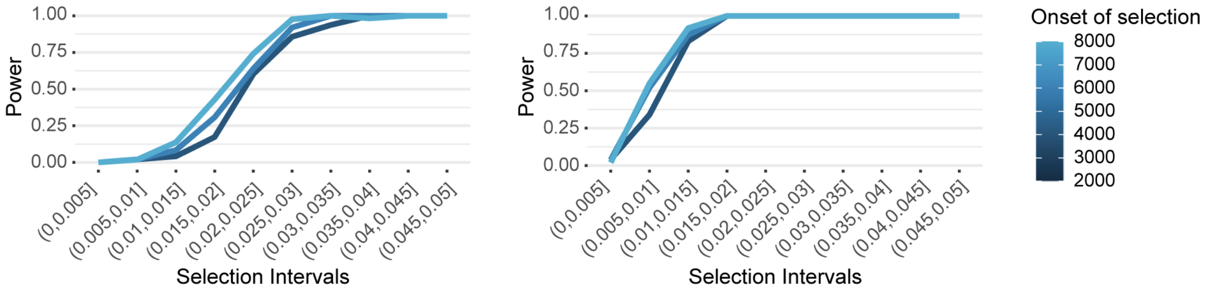


**Figure S2. Power to detect selection as a function of the time of selection onset**
Power in the case of positive selection for an allele frequency distribution matching real data for variants with frequencies between 0 and 0.2 (left panel) or between 0.2 and 0.6 (right panel), for an empirical significance level of $10^{-5}$. Power estimates for each combination of selection coefficient and time of selection onset are based on 100 selection coefficient estimations.

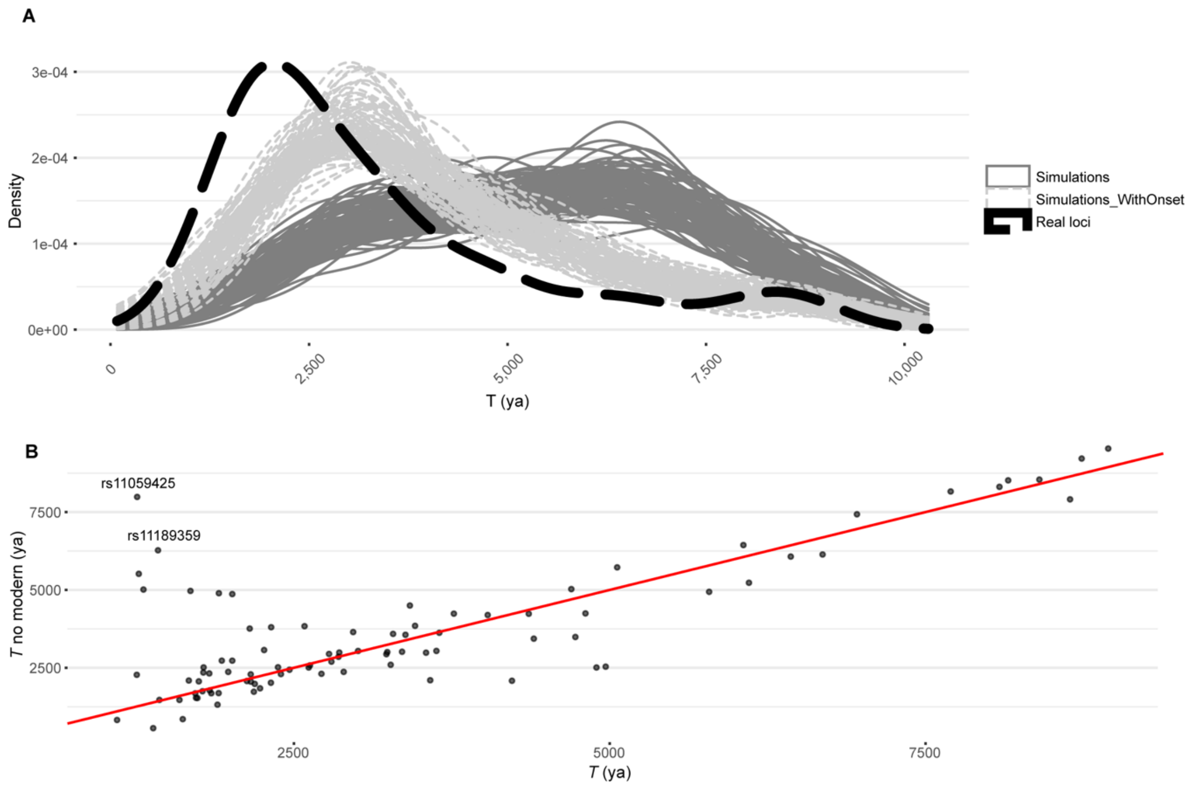


**Figure S3. Distribution of the time of selection onset for the variant with the smallest $p_{\mathrm{sel}}$ at each of the 89 candidate positively selected loci**

(A) Densities are shown for the distribution of selection onset estimates (*T*) for the variant with the smallest $p_{\mathrm{sel}}$ at each of the 89 positively selected loci or the *T*s of 100 sets of 89 independent simulated variants matching the allele frequencies and selection strengths of each of the 89 variants (continuous dark gray lines), or 100 sets of 89 independent simulated variants matching the allele frequencies, selection strengths and times of selection onset of the 89 variants (continuous light gray lines).

(B) *T* estimates for the 89 positively selected variants with minimal *p* values at each of the 89 loci determined with (*x*-axis) or without (*y*-axis) the use of modern DNA.

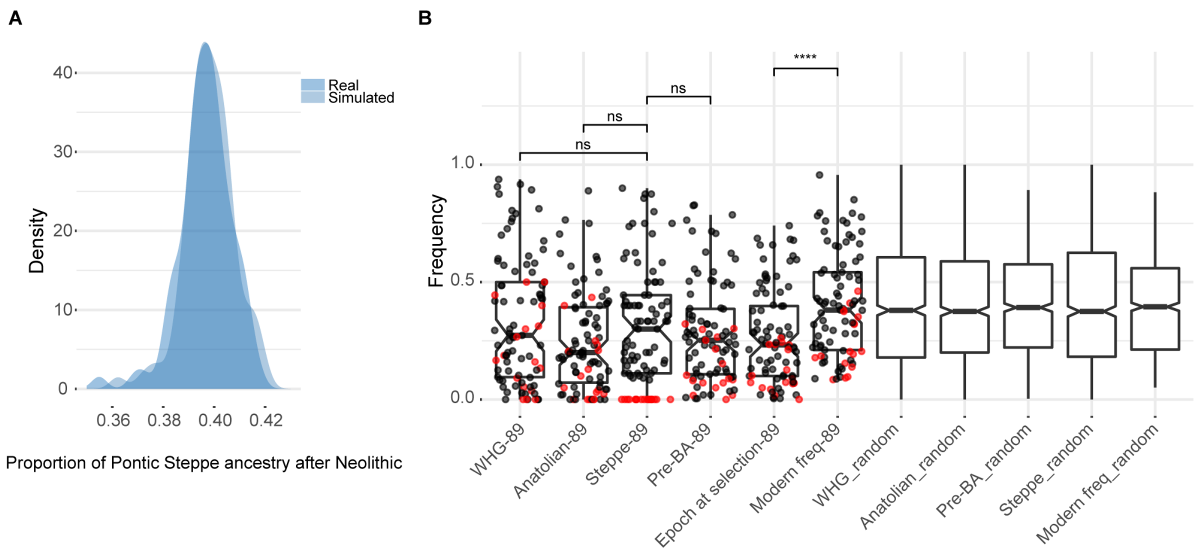


**Figure S4. Adaptive admixture has not been the main driver of positive selection in post-Neolithic Europe**

(A) Distribution of the estimated mean proportion of Pontic Steppe ancestry for the post-Neolithic carriers of the allele with smallest $p_{\text{sel}}$ at each of the 89 candidate positively selected loci (darker blue, 'Real'), or for the simulated aDNA used to obtain $s$ and $T$ estimates for each of the 89 variants (lighter blue, 'Simulated').

(B) Frequencies for the 89 alleles with smallest $p_{\text{sel}}$ at each of the 89 candidate positively selected loci (boxplots with dots) or for 1,000 random variants matched on DAF (boxplots without dots). Frequency distributions are represented for different ancestry groups (Western hunter gatherer [WHG], Anatolian and Steppe) and differences between these distributions were assessed with a Wilcoxon test (not significant [ns] for all three tests). Frequencies are also represented for different epochs (pre-Bronze Age samples [Pre-BA], samples from the epoch in which selection is estimated to have begun [Epoch at selection] and modern samples [Modern Freq]). A Wilcoxon test was also used to test for differences in frequency distributions between "Epoch at selection-89" and "Modern Freq-89". This difference was significant, as expected. The red dots indicate alleles with a frequency estimation of 0 among individuals with Pontic-Steppe ancestry. Individuals were defined as belonging to a given ancestry category if their proportion of the corresponding ancestry was >90%. The Steppe category was also limited to individuals dating back more than 4,500 years, the Anatolian category was limited to individuals dating back more than 7,500 years, and the WHG category was limited to individuals dating back more than 8,000 years. Frequency trajectories for these 89 alleles can be found in **Table S2**.

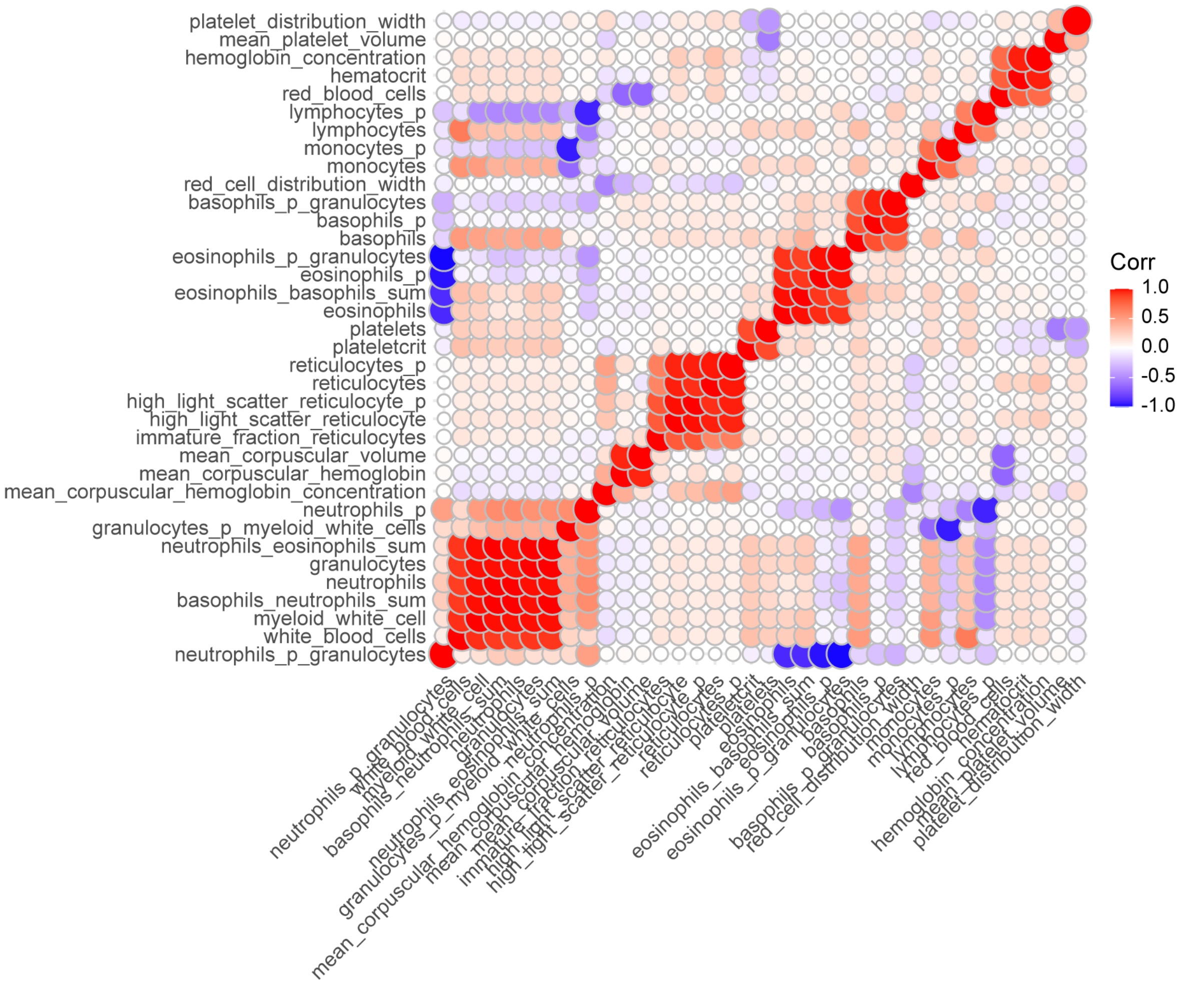


**Figure S5. Genetic correlations between the study hematopoietic traits**
Genetic correlations were computed using ‘ldsc’ (**STAR Methods**).

**Table S1. Evolutionary parameters for previously published positively selected loci**

| CHR | Lead SNP[a] | POS hg19 | DA | *S*[b] | T (ya)[b] | Genes[c] | $p_{sel}$ |
|---|---|---|---|---|---|---|---|
| 2 | *rs4988235* | 135.3-137.3 | A | 0.082 (0.087) | 6,102 (3,401) | *MCM6, LCT* | 0 |
| 5 | *rs185146* | 33.8-34 | T | 0.059 (0.061) | 4,226 (12,583) | *SLC45A2* | 0 |
| 6 | *rs3130673* | 29.9-33.1 | T | 0.077 (0.067) | 8,401 (10,901) | *MHC region* | 0 |
| 11 | *rs174537* | 61.5-61.6 | G | 0.013 (0.013) | 6,685 (10,263) | *FADS1, FADS2* | $2.7 \times 10^{-5}$ |
| 4 | *rs13149231* | 38.7-38.8 | T | 0.016 (0.020) | 6,570 (11,300) | *TLR1* | $5.8 \times 10^{-5}$ |
| 12 | *rs11065987* | 111.9-112.6 | G | 0.013 (0.028) | 5,597 (6,456) | *ATXN2, SH2B3* | $5.2 \times 10^{-4}$ |
| 11 | *rs1540129* | 71.1-71.2 | C | 0.020 (0.021) | 3,333 (3,251) | *DHCR7* | 0 |
| 11 | *rs10765770* | 88.5-88.9 | A | 0.019 (0.021) | 6,058 (8,618) | *GRM5* | $2.0 \times 10^{-6}$ |
| 5 | *rs27879* | 131.4-131.8 | A | 0.020 (0.027) | 8,884 (11,957) | *SLC22A4* | $1.9 \times 10^{-5}$ |
| 6 | *rs9468413* | 28.3-28.7 | C | 0.049 (0.042) | 6,057 (8,927) | *ZKSCAN3, ZSCAN31* | 0 |
| 13 | *rs9566230* | 38.1-38.8 | C | 0.017 (0.029) | 1,530 (26,076) | - | $3.8 \times 10^{-5}$ |
| 11 | *rs72878978* | 29.9-30.1 | A | 0.016 (0.011) | 2,387 (1,865) | *HERC2, OCA2* | $2.9 \times 10^{-3}$ |

[a]The SNP with the smallest $p_{sel}$ at each of the 12 previously published positively selected loci.[43]
[b]*T* and *s* values are shown for estimates based on $T \sim$ U[-10,000-0] ya or in parentheses for $T \sim$ U[-30,000-0] ya, as priors used in the simulations.
[c]Most relevant gene(s) for the lead SNP of each locus.

**Table S2. The 89 variants with the smallest $p_{sel}$ at each of the 89 candidate positively selected loci**
(see separate Excel file)

**Table S3. Variants underlying the evolutionary history of the ABO locus**

| **SNP (ABO type)** | **In 1240k array** | **Proxy (allele)** | **Correlated alleles - $r^2$ (ABO type)** | **$p_{sel}$ (negative or positive)** | **Disease association (direction)** |
|---|---|---|---|---|---|
| *rs8176719>TC* (non-O type) | No | No | - | - | Malaria (susceptibility) |
| *rs8176746>G* (B type) | No | No | - | - | - |
| *rs41302905>T* (O type) | Yes | No | - | 0.04 (negative) | - |
| *rs505922>C* | Yes | Yes (rs8176719) | C=TC - 0.87 (non-O type) | 0.01 (positive) | - |
| *rs8176749>T* | Yes | Yes (rs8176746) | T=G - 1 (*B* type) | $1.2 \times 10^{-4}$ (positive) | - |
| *rs8176743>T* | Yes | Yes (rs8176746) | T=G - 1 (B type) | 0.05 (positive) | - |
| *rs9411378>A* | No | Yes (rs8176719) | A=TC - 0.64 | - | COVID-19 (susceptibility) |
| *rs635634>C* | Yes | Yes (rs8176719) | C=TC - 0.4 | $6.4 \times 10^{-4}$ (positive) | TS and CIE (protection) |

Note: TS stands for tonsillectomy and CIE for childhood ear infection. $r^2$ are given for EUR populations of 1KG. We found no variant correlated with rs9411378 with an $r^2$ >0.8 in the 1240k array.

**Table S4. Lead GWAS SNPs of all infectious and autoimmune traits**
(see separate Excel file)

**Table S5. Top SNPs underlying the increase in the risk of inflammatory disorders over time**

| SNP | Phenotype | Consequence | *p* (PRS)[a] | *p* (eQTL) | Gene[b] |
|---|---|---|---|---|---|
| *rs2188962* | CD | Intronic | $7.04 \times 10^{-19}$ | $1.8 \times 10^{-20}$ | *IRF1* |
| *rs2188962* | IBD | Intronic | $5.09 \times 10^{-19}$ | $1.8 \times 10^{-20}$ | *IRF1* |
| *rs11066188* | CD | Upstream | $2.82 \times 10^{-12}$ | $1.2 \times 10^{-49}$ | *SH2B3* |
| *rs11066188* | IBD | Upstream | $1.80 \times 10^{-11}$ | $1.2 \times 10^{-49}$ | *SH2B3* |
| *rs492602* | CD | Synonymous | $2.08 \times 10^{-10}$ | - | *FUT2* |
| *rs1456896* | IBD | Intergenic | $2.25 \times 10^{-4}$ | $1.6 \times 10^{-3}$ | *IKZF1* |
| *rs10774679* | COVID | Upstream | $2.05 \times 10^{-4}$ | - | *OAS1* |

[a]*p* values were obtained after adjustment for ancestry (factor components) and sample location (longitude and latitude)
[b]Either the gene eQTL/sQTL (rs2188962, rs11066188, rs1456896 and rs10774679) or the gene in which the variant occurs (rs492602)

**Table S6. Candidate negatively selected missense variants at conserved positions**
(see separate Excel file)

**Table S7. List of ancient samples used in this work**
(see separate Excel file)

**Table S8. Filtering bias for variants close to indels or not in the capture dataset**

| SNP | Shotgun_c[a] Capture_na | Shotgun_nc[a] Capture_na | Capture[b] frequency | Shotgun[b] frequency | 1KG frequency | Nearby indel |
|---|---|---|---|---|---|---|
| *rs264272* | 0.72 | 0.014 | 0.12 | 0.49 | 0.51 | Yes |
| *rs2792600* | 0.78 | 0.051 | 0.05 | 0.38 | 0.45 | Yes |
| *rs4283567* | 0.54 | 0.027 | 0.19 | 0.41 | 0.49 | Yes |
| *rs8076492* | 0.61 | 0.054 | 0.28 | 0.56 | 0.56 | Yes |
| *rs13225222* | 0.57 | 0.025 | 0.04 | 0.14 | 0.19 | Yes |
| *rs7248807* | 0.73 | 0.022 | 0.07 | 0.26 | 0.39 | Yes |
| *rs11621931* | 0.67 | 0.024 | 0.01 | 0.08 | 0.10 | Yes |
| *rs13126794* | 0.71 | 0.030 | 0.02 | 0.15 | 0.17 | Yes |
| *rs10763588* | 0.59 | 0.046 | 0.20 | 0.54 | 0.56 | Yes |
| *rs996759* | 0.041 | 0.026 | 0.42 | 0.28 | 0.45 | No |
| *rs13180583* | 0.052 | 0.029 | 0.40 | 0.37 | 0.39 | No |
| *rs6437191* | 0.079 | 0.017 | 0.40 | 0.44 | 0.41 | No |
| *rs16961871* | 0.034 | 0.029 | 0.13 | 0.15 | 0.20 | No |
| *rs12615153* | 0.026 | 0.025 | 0.56 | 0.60 | 0.67 | No |
| *rs2835900* | 0.037 | 0.034 | 0.27 | 0.29 | 0.35 | No |
| *rs10146186* | 0 | 0.024 | 0.07 | 0.06 | 0.06 | No |
| *rs11592504* | 0.11 | 0.094 | 0.07 | 0.05 | 0.10 | No |
| *rs72649559* | 0.018 | 0.026 | 0.16 | 0.14 | 0.19 | No |

Note: Columns 2 and 3 indicate the proportion of individuals with a status in the shotgun database but absent from the capture database.
[a]'_c' stands for carriers, '_nc' stands for non-carrier and '_na' stands for absent
[b]Frequencies were obtained by averaging frequencies across all past epochs

**Table S9. Primers used for experiments on *LBP***
(see separate excel file)
